# Somatic mobility of transposons is explosive and shaped by distinct integration biases in *Arabidopsis thaliana*

**DOI:** 10.1101/2025.07.14.664700

**Authors:** Heena Ambreen, Basile Leduque, Leandro Quadrana, R. Keith Slotkin, Alexandros Bousios, Hans-Wilhelm Nützmann

**Author notes:** **Author’s Email addresses:** Heena Ambreen. Hans-Wilhelm Nützmann.

## Abstract

**Background:** The movement of transposable elements (TEs) in somatic cells generates genetic and phenotypic diversity across the soma of individual organisms. This process is especially important in plants that display a sessile lifestyle, can regenerate from somatic cells, and continuously establish their germline from somatic stem cells. Yet, the scale and integration biases by which TEs may shape the plant soma remain unknown.

**Results:** We adapt high-throughput TE sequencing and introduce a new computational pipeline, “deNOVOEnrich”, to identify and analyse somatic transposition events in *Arabidopsis thaliana*. We detect ∼200,000 new TE insertions across wild-type and epigenetic mutant backgrounds, revealing en masse active transposition in the plant soma. Similar to germline dynamics, somatic integration is non-random and TE-specific. Focusing on the ONSEN, EVADE and AtCOPIA21 families, we find they preferentially integrate into chromosomal arms and genic DNA, and are enriched over H2A.Z, H3K27me3 and H3K4me1 labelled chromatin. However, ONSEN dynamically targets active and open chromatin, while EVADE displays a depletion in these regions, consistent with niche targeting at the local epigenetic level and spatial distribution across the genome ecosystem. We further demonstrate that environmentally-responsive genes such as resistance (R) genes and biosynthetic gene clusters are preferred integration targets, highlighting the adaptive potential of somatic transposition.

**Conclusions:** Our work demonstrates that transposon amplification is an explosive and structured event that generates cellular mosaicism in the soma of *A. thaliana*. We speculate that this phenomenon is common across plants under permissive conditions, both in experimental and natural settings.

## Background

Transposable elements (TEs) typically occupy large fractions of eukaryotic genomes, and are crucial for generating novel genetic and phenotypic variation [Anderson et al., 2019; Akakpo et al., 2020; Wells and Feschotte, 2020]. The major contribution of TEs in shaping genome structure and function is well-documented across host taxa [Lisch, 2009; Bourque et al., 2018], however, much of our understanding of the genome-wide impact of TEs has been shaped by studies focussing on heritable transposition events. In contrast, research on the large-scale dynamics of somatic TEs has not been on par with exploring insertions of TEs in the host germline [Bourque et al., 2018], particularly in plant species. One possible explanation is the long-held idea of complete suppression of TE activity in somatic cells [Haig, 2016; Loreto & Periera, 2017]. Furthermore, new transposon insertions in the soma are typically not inherited through generations, and are thus considered to have limited evolutionary relevance for both the host and the TE [Kazazian, 2011; Loreto & Pereira, 2017]. In addition, somatic TE insertions are rare variants limited to single cells or small cell populations, and, hence, their detection is technically daunting and a plausible reason for underestimating levels of somatic TE transposition [Treiber & Waddell, 2017; Siudeja et al., 2021].

Several studies have provided evidence for frequent TE movement in somatic cells [Baillie et al., 2011; Siudeja et al., 2021; Mendez-Dorantes & Burns, 2023]. In humans and mice, LINE-1 elements transpose extensively in brain cells, introducing non-heritable changes in neuronal somatic genomes that contribute to cellular heterogeneity and may potentially influence cognitive functions [Baillie et al., 2011; Bodea et al., 2024]. Similar release of high transposition activity has been observed in cancerous and aged somatic cells in humans [Lee et al., 2012; Rodriguez-Martin et al., 2020; Gorbunova et al., 2021]. In *Drosophila melanogaster*, regular TE movement has been characterized in brain and gut tissues, as well as in aged individuals, demonstrating that somatic mosaicism mediated by TE insertional mutagenesis is an ongoing genomic process [Chang et al., 2019; Siudeja et al., 2021; Yang et al., 2022]. Outside these model metazoans, TE mobilization in somatic tissues is poorly characterised at genome-wide scales.

In plants, the implications of somatic transposition are more far-reaching than in animals. Unlike animals, plant cell differentiation is not dependent solely on cell division lineage patterns, and cells are not set aside early in development for the germline; instead, plants establish their germline in late adulthood from stem cells. In addition, upon wounding or induced in the laboratory, any somatic plant cell has the ability to redifferentiate and contribute to the production of germ cells. This suggests that TE insertions in somatic genomes have higher likelihood of transmitting into the progeny and become a source of novel genetic variability [Bourque et al., 2018]. Yet, most studies have focussed on forward genetics approaches to explain phenotypes showing somatic variegation caused by random TE insertions [Coen et al., 1986; Alleman and Kermicle, 1993; Iida et al., 2004; Xu et al., 2010; Zhang et al., 2024] or detected clonal diversity in vegetatively propagated crop species that is derived from TE transposition [Kobayashi et al., 2004; Fernandez et al., 2010; Carrier et al., 2012; Wang et al., 2021b; Ban et al., 2022].

In maize, studies have shown that somatic transposition leads to differentially colored sectors in kernels [McClintock, 1984, McCarty et al., 2013], while stress-induced transcriptional activation of TEs in somatic cells is well documented in plants [Ito et al., 2011, Makarevitch et al., 2015]. These studies are consistent with TEs being active in the plant soma and important for phenotypic somatic diversity and plant stress physiology. However, the large-scale dynamics of somatic transposition are poorly characterised in plants, leaving a significant gap in our understanding of genome-wide rates of somatic TE proliferation and their impact on genome stability, function and evolution. In addition, investigating somatic transposition in-depth provides a unique opportunity to capture *de novo* integration patterns of TEs before selection removes deleterious insertions [Sultana et al., 2019; Cao et al., 2023], which is a fundamental aspect of TE and genome biology.

Here, using the well-characterized TE mobilome in *Arabidopsis thaliana*, we set out to reliably assess the large-scale dynamics of somatic transposition in a plant species, and to determine the *de novo* integration preferences of actively transposing TEs. To release TE suppression and trigger high TE activity in our model system, we employ heat stress and epigenetic mutants. We then apply transposable element display sequencing (TEd-seq) [Vendrell-Mir et al., 2025] to identify rare TE-associated fragments from targeted genomic libraries, and design a new computational pipeline, “*deNOVOEnrich*” to detect and characterize somatic TE insertions to base pair resolution. This set-up enables us to identify over 200,000 new transposition events for three exemplar LTR retrotransposon families. Analysis of their integration sites reveals shared and unique genomic and epigenomic target site preferences, and a distinct preference for environmentally-responsive genes such as resistance (R) genes and biosynthetic gene clusters. Overall, we provide evidence that TEs mobilize en masse in somatic cells of *A. thaliana* plants, opening the field for future studies on the functional impact of somatic genome plasticity across species.

## Results

### Monitoring active TEs in Arabidopsis seedlings

An essential requirement to study somatic insertions is the identification of TE families that can retain high levels of activity and generate large numbers of integration events. To boost transposition in *A. thaliana*, we used four epigenetic mutant lines that are known to compromise TE silencing at both the transcriptional and posttranscriptional levels, i) the methyltransferase 1 (*met1*) mutant that leads to loss of maintenance of cytosine methylation in the CG sequence context [Kankel et al., 2003], ii) the plant-specific RNA polymerase IV (*polIV*) mutant that is crucial for small interfering RNA (siRNA) production during the upstream phase of RNA-directed DNA methylation (RdDM) [Kanno et al., 2005], iii) the plant-specific RNA polymerase V (*polV*) mutant that produces the scaffold transcript of the TE to anchor the RdDM machinery to chromatin [Böhmdorfer et al., 2016; Sigman et al., 2021] and iv) the *polIVpolV* double mutant (**Additional file 2: Table S1**). We examined several TE families that are known to be “epigenetically-induced” in these *A. thaliana* mutants [Tsukahara et al., 2009, Quadrana et al., 2019; Lee et al., 2020]: the LTR retrotransposon families EVADE (AtCOPIA93), AtCOPIA21, AtCOPIA63, AtCOPIA51, AtCOPIA31, ATGP3 and AtCOPIA52 (SISYPHUS). We also applied heat stress to our plants, because of the presence of heat-responsive TEs, namely the LTR retrotransposon ONSEN (AtCOPIA78) family that has been thoroughly studied as a model TE for stress-mediated activation in plants [Ito et al., 2011; Cavrak et al., 2014; Gaubert et al., 2017]. Our set of TE families contain between one and eleven intact elements in the *A. thaliana* ColCEN reference genome, and overall occupy 0.16% of genomic DNA (**Additional file 2: Table S2**).

Our current understanding of ONSEN heat-induction is limited to an application period of 24-48 hours [Ito et al., 2011; Cavrak et al., 2014; Thieme et al., 2022]. To better discern when maximal transcription is achieved, we profiled ONSEN transcript levels with qRT-PCR in a wider range of heat exposure and treatments (**Figure 1A-C; Additional file 1: Fig. S1;** see Methods). In leaf tissues of wild-type plants, ONSEN transcription showed a continuous increase for seven days of heat exposure (37°C), beyond which transcript levels fell and seedlings died **(Figure 1A,B; Additional file 1: Fig. S2A)**. Cold priming did not yield any significant effect on transcript abundance (**Additional file 1: Fig. S1**), while sampling after a 48-hour recovery phase decreased ONSEN transcript levels in relation to sampling directly after the heat-stress **(Figure 1B)**. The four mutant lines followed a similar trend as wild-type plants, whereby ONSEN transcription correlated strongly with the duration of applied heat (**Figure 1A, C**). However, all lines showed adverse growth damage after three days, with symptoms being most severe in *polIVpolV* mutant plants (**Additional file 1: Fig. S2B,C**). Among the other TE families, we could detect transcripts based on the qRT-PCR profiling only for EVADE and AtCOPIA21 in the epigenetic mutant lines, with *met1* showing the highest activation (**Figure 1D**).

**Figure 1.**
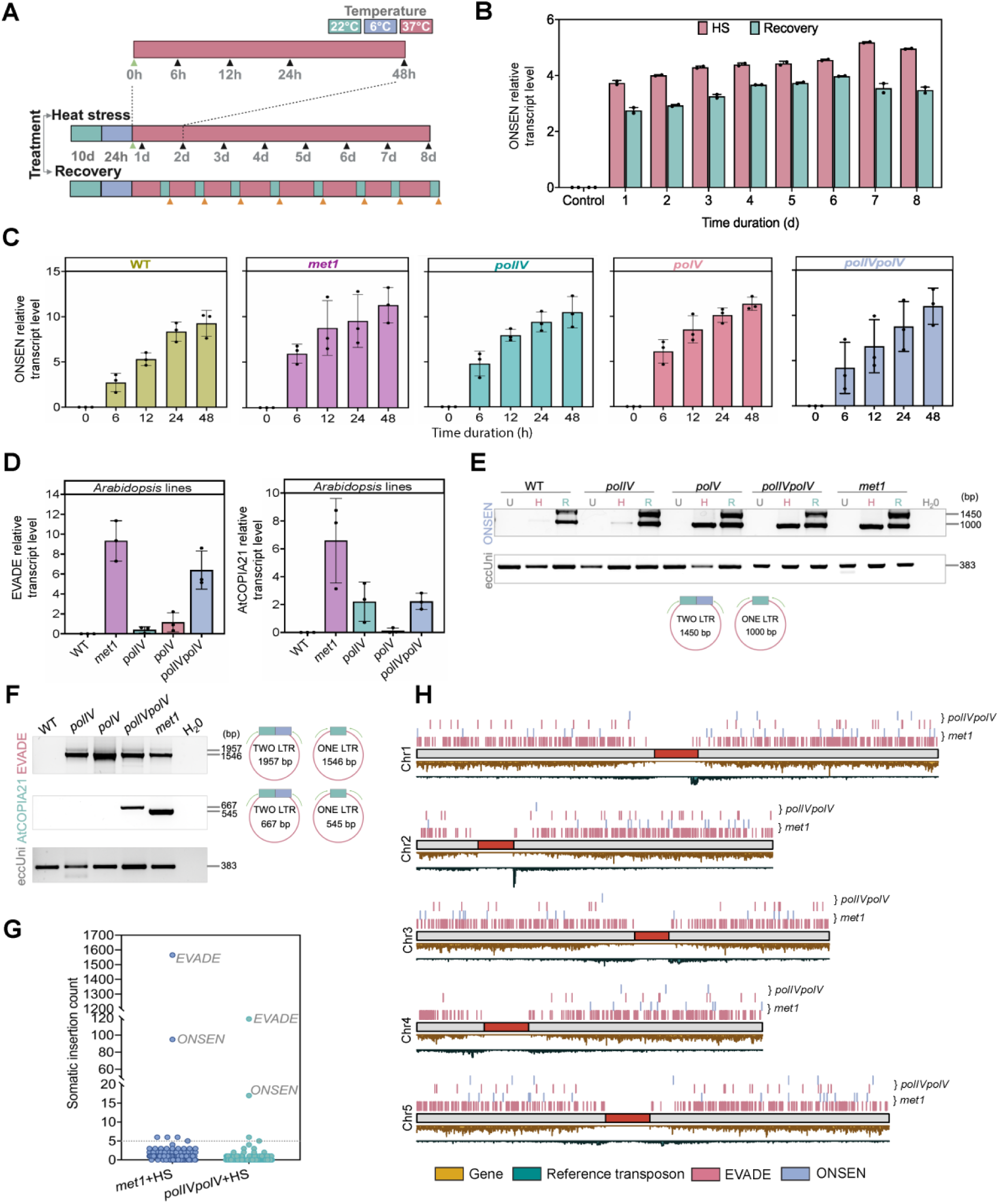
**Activation of TE families in *A. thaliana* seedlings**. **A.** Heat treatments and timepoints for assessing ONSEN transcription levels with qRT-PCR. Sampling timepoints are indicated with green (control), black (heat stress), and orange (recovery) arrowheads. Recovery samples were taken after incubation for 48 hours at 22°C post the heat stress. **B**. ONSEN qRT-PCR expression profiles under 8-days heat regime (n=2). Leaf tissues from ∼100 seedlings were pooled for each replicate. **C.** ONSEN qRT-PCR expression profiles at regular intervals extending up to 48 hrs in wild-type (WT) and mutants (n=3). **D.** Relative expression levels of EVADE and AtCOPIA21 (n=3) in unstressed tissue of *Arabidopsis* lines. Mean ± SD plotted for qRT-PCR analysis in B-D. **E, F.** Inverse PCR profiles for detection of eccDNA for ONSEN **(E)** EVADE and AtCOPIA21 **(F)**. Unstressed (U), 48h heat-stressed (H) and 48h recovery post HS (R) samples were used for ONSEN. Unstressed plant material was used for EVADE and AtCOPIA21. Expected size for two and single LTR products have been indicated. eccDNA molecule (eccUni) universally observed in *Arabidopsis* tissues (Wang et al., 2021a) has been used as positive control. **G.** Number of *de novo* somatic insertions of different TE families identified in *met1* and *polIVpolV* heat-stressed leaf tissues using whole genome re-sequencing. **H.** Genome-wide distribution of somatic ONSEN and EVADE insertions shown in G. Gene and reference TE densities are shown in 10 kbp windows. Centromere location has been shown by red blocks.

As a more direct marker of TE mobility, we detected TE extrachromosomal circular DNA (eccDNA) that is generated after reverse transcription [Zhang et al., 2023]. We examined the formation of eccDNA by inverse PCR profiling that captures LTR junctions of eccDNA molecules. Our analysis of seedlings exposed to a 48-hour heat-stress with and without a 48-hour recovery phase revealed the presence of two abundant forms of ONSEN eccDNA with one or two LTRs in all lines (**Figure 1E**). The two-LTR eccDNA product was prominent in the recovery phase, consistent with extrachromosomal DNA copies accumulating in the cytoplasm after the synthesis of excessive levels of TE transcripts, or with integration of new copies slowing down after the heat-stress window. EVADE eccDNA was abundant across all mutant lines and mostly contained one LTR, while AtCOPIA21 eccDNA was detected only in the *met1* and *polIVpolV* double mutant (**Figure 1F**). Identity of the variable LTR forms of ONSEN and EVADE eccDNA was confirmed by Sanger sequencing (**Additional file 1: Fig. S3**). We also found varying levels of eccDNA for more families in our mutant lines but not the wild-type controls (**Additional file 1: Fig. S4**), indicating distinct interactions between TEs and the epigenetic silencing pathways. Collectively, our assessment of known environmentally and epigenetically-induced TE families using qRT-PCR and eccDNA analyses points towards broad activation of TEs, with ONSEN, EVADE and AtCOPIA21 showing the highest levels of activity.

### Whole genome sequencing supports somatic transposition of ONSEN and EVADE

To explore whether high levels of transcription and eccDNA can lead to transposition in somatic cells, we performed high coverage (>100x fold) Illumina short-read whole-genome sequencing on 48 hour heat-stressed leaf tissues pooled from ∼100 seedlings of *met1* and *polIVpolV,* the two mutant backgrounds where most TE families show evidence of high activity (**Figure 1A-F, Additional file 1: Fig. S4**). We mapped *de novo* somatic insertions with the TEMP2 software [Yu et al., 2021], which uses split and discordant reads for the identification of novel insertions. Each identified insertion was allocated to a specific family using the curated TE library of the Col-CEN reference genome. We detected 1,803 and 198 putative insertions in *met1* and *polIVpolV*, respectively, with seven families substantially contributing to transposition load by having ≥5 insertions (**Figure 1G**) (**Additional Data 3**). ONSEN and EVADE exhibited the highest levels of mobility with 112 (5.6%) and 1,684 (84.2%) insertions for both mutants combined (**Figure 1G**). The majority of these insertions were localized on the chromosomal arms (**Figure 1H**).

### Explosive bursts of somatic transposition detected by targeted sequencing

While high coverage whole-genome sequencing can provide evidence for somatic TE mobilization, the inherent limited sensitivity of a genome-wide approach restricts our ability to assess the true extent of integration in the somatic pool of cells. We therefore leveraged a recently developed TE-enrichment sequencing approach, TEd-seq, that is designed to identify non-reference low frequency insertions for single TE families with ultra sensitivity and specificity in large numbers of individuals, such as in “evolve and resequence” population experiments [Vendrell-Mir et al., 2025; Tsukahara et al., 2025] (**Figure 2A**). To adapt the bioinformatic analysis to the expected lower frequency of TE insertions within a DNA sample of somatic tissue, we developed a new computational pipeline, termed “*deNOVOEnrich*”. This pipeline uses deeply-filtered, high-confidence split reads to map the *de novo* insertional landscape to base pair resolution (**Figure 2A, Additional file 1: Fig. S5**) (see Methods). *deNOVOEnrich* applies read coverage to differentiate single-generation somatic (≤5 reads) from non-reference segregating insertions (≥30), with the latter occurring, for example, during seed propagation cycles prior to TEd-seq application (**Figure 2A,B, Additional file 1: Fig. S5**).

**Figure 2.**
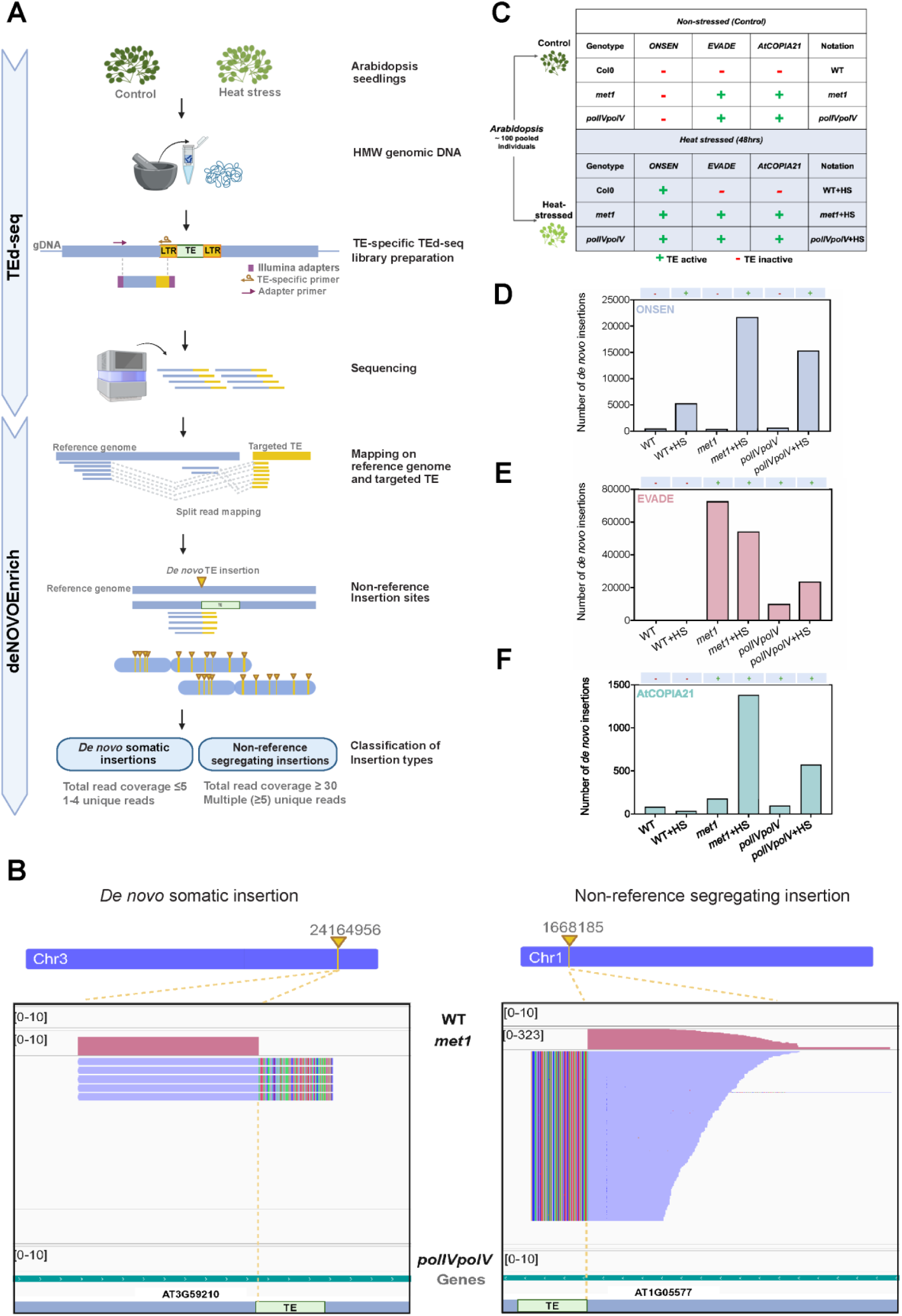
**Somatic transposition of ONSEN, EVADE and AtCOPIA21 in *A. thaliana* leaves**. **A.** Workflow for the precise detection of *de novo* somatic and non-reference segregating insertions. *TEd-seq*: Targeted sequencing libraries are generated for specific TEs and sequenced at high coverage with Illumina. *deNOVOEnrich:* New insertion sites are identified based on split-reads, whereby reads that partially map to the reference genome, and their unmapped region shows strong homology to the TE end, are used to identify genome-wide transposition events. The insertions are classified as somatic or non-reference segregating insertions based on coverage thresholds (see Methods and **Additional file 1: Fig. S5** for step-by-step details on the *deNOVOEnrich* pipeline). **B.** Genome browser view of representative *de novo* somatic (left) and non-reference segregating (right) insertions of EVADE in the *met1* mutant. Read coverage (pink) and alignment (blue) tracks are shown for the TE-genome junction of the new insertions. Clipped read segments corresponding to the TE end are indicated by rainbow-colored portions. Tracks for the same genomic location in wild-type and *polIVpolV* are included. Chromosomal position (top) and the specific insertion coordinates within two different genes (bottom) are shown in the ideograms. (for information on non-reference segregating insertions identified in our analysis see **Additional file 1: Supplementary Information 1** and **Fig. S15**) **C.** Experimental conditions used for TEd-seq. The expected activity status (“+” for active, “-” for inactive) is shown for each combination of genome background and TE family. **D-F.** Number of somatic TE insertions for ONSEN (D), EVADE (E) and AtCOPIA21 (F) in experimental combinations of (C). The expected activity status is indicated above each plot.

Based on our initial screenings for active TE families (**Figure 1**), we generated TEd-seq data for ONSEN, EVADE and AtCOPIA21 in wild-type, *met1* and *polIVpolV* mutant plants under stress (37°C for 48 hours) and non-stress conditions (**Figure 2C**). These combinations generate predictions about the activation patterns of each family, which then serve as controls to estimate false discovery rates (FDR) and enhance the robustness of the analysis. For example, ONSEN is not anticipated to transpose in the absence of heat-stress in either wild-type or mutant plants, EVADE should only transpose in mutant lines regardless of the application of heat, and no family should transpose in wild-type with no heat-stress (**Figure 2C**). Across all TEd-seq libraries, we generated 132.5 Gbp of data using 2×150 bp paired-end reads, which ranged between 8.8 to 32.6 Gbp per library and was derived from leaves of a pool of ∼100 seedlings (**Additional file 2: Table S3**).

After normalising for library size, we detected for ONSEN a total of 5,310, 15,355 and 21,731 somatic insertions in heat-stressed wild-type, *polIVpolV* and *met1* plants compared to 473, 639 and 432 in non-stress conditions of the same lines (**Figure 2D, Additional file 1: Fig. S6**). This represents a 11-, 24-and 50-fold increase and is consistent with low FDRs in our pipeline. Notably, ONSEN showed significantly higher levels of transposition in the mutant lines, indicating enhanced propagation in epigenetically repressed conditions in addition to the effect of heat. For EVADE, loss of epigenetic control resulted in massive propagation in somatic cells of non-stressed *polIVpolV* and *met1* plants with 10,022 and 72,678 new insertions respectively, which are 50 and 363 times higher than the FDR levels observed in non-stressed wild-type (200 insertions) **(Figure 2E, Additional file 1: Fig. S7)**. Application of heat in the mutant plants did not significantly alter transposition rates (1.3-2 fold difference; 23,621 insertions in *polIVpolV* and 54,235 in *met1*) (**Figure 2E, Additional file 1: Fig. S7**). In contrast to ONSEN and EVADE, AtCOPIA21 displayed less intense transposition in the mutant lines (**Figure 2F, Additional file 1: Fig. S8**). In fact, the most notable increase was observed in the heat-stressed mutant plants, which showed >5 fold difference in relation to wild-type (572 and 1,384 in *polIVpolV* and *met1*, respectively). Taken together, our TEd-seq/deNOVOEnrich analysis has captured hundreds of thousands of new insertions in the Arabidopsis soma, establishing that somatic transposition is an explosive ongoing event under conditions of weakened epigenetic control or environmental stress.

### Genomic features influencing *de novo* somatic integration

Analysis of TE insertion patterns often show non-random distribution along chromosomes, which can be attributed to the combined effect of integration preferences and post-integration selection acting on each site [Sultana et al., 2017; Cao et al., 2023]. In plants, TE integration patterns have been mostly studied by analysing chromosomal distribution of TEs in generations subsequent of TE amplification (i.e. post-selection), which obscures true insertion biases. Our investigation, however, of genome-wide *de novo* somatic transposition within a single generation provides a unique opportunity to capture integration patterns before selection alters their distribution.

We first explored the distribution of somatic insertions in the three main compartments of the *A. thaliana* genome, the centromeres (9.6% of genomic DNA), pericentromeres (12.1%), and chromosomal arms (78.3%). Centromere boundaries were defined in the Col-CEN reference assembly [Naish et al., 2021], while the junctions between the pericentromeres and the chromosomal arms were recently resolved based on mapping thousands of meiotic recombination crossovers [Fernandes et al, 2024]. Reflecting a strong affinity for euchromatin, all three families, and especially EVADE (94.4% of insertions), showed a universal preference to integrate into chromosome arms (Permutation test; p<0.001) (**Figure 3A-D; Additional file 1: Fig. S9A**), while being underrepresented in pericentromeres and absent from centromeres that are only invaded by ATHILA LTR retrotransposons [Naish et al., 2021; Wlodzimierz et al., 2023] (**Figure 3A-D; Additional file 1: Fig. S9A**).

**Figure 3.**
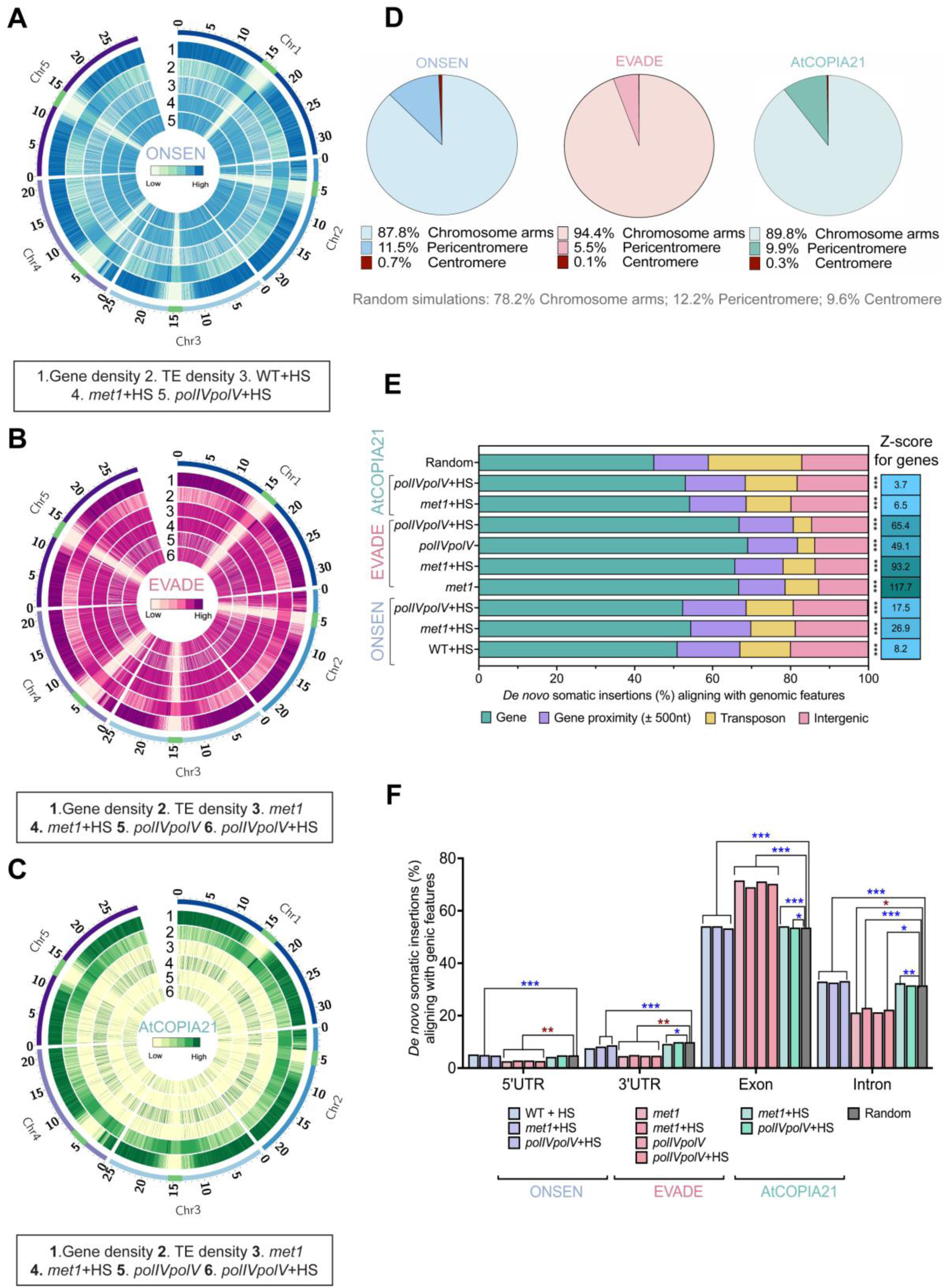
**Topology of somatic insertions in the *A. thaliana* genome**. **A-C**. Circos plots showing somatic insertions along the *A. thaliana* chromosomes for ONSEN (**A**), EVADE (**B**), and AtCOPIA21 (**C**). Green blocks in outermost ribbons indicate centromere positions. Gene and TE density have been calculated in 20kb non-overlapping windows. **D.** Proportion of somatic insertions in chromosomal arms, pericentromeres and centromeres. To calculate the expected frequency of insertion in the three main genomic compartments, simulations were conducted 1000 times with permutation test. **E**. Fraction of somatic insertions in genes, 500bp windows proximal to genes, reference TEs and intergenic regions. Permutation tests were applied to evaluate the level of significance for each combination of genome background and TE family (***p<0.001; Z scores are shown for genes). Z scores for other genomic regions are provided in **Additional file 1: Fig. S9B**. For AtCOPIA21, only heat-stressed samples of mutant lines were analysed, following the number of somatic insertions detected as shown in Figure 2F. **F.** Fraction of somatic insertions in different genic components. Permutation tests were applied to evaluate the level of significance for genic components and somatic insertions (***p<0.001, **p<0.01, *p<0.05). Positive association marked by blue asterisk while negative association shown by red asterisk. Only statistically significant relationships have been indicated. Permutation Z scores for genic regions are provided in **Additional file 1: Fig. S9C**.

We further partitioned the genome into (i) genes, (ii) 500bp regions flanking genes, (iii) reference TEs, and (iv) intergenic space to associate somatic integration with underlying genomic features at the regional scale. Among the three families, and following the strong bias for euchromatin, EVADE was significantly enriched in genes (67% on average across all mutants, Permutation test; p<0.001) (**Figure 3E**;

**Additional file 1: Fig. S9B**), and, in particular, exonic DNA (Permutation test; p<0.001) (**Figure 3F; Additional file 1: Fig. S9C**), while being substantially less present within the TE space (**Figure 3E; Additional file 1: Fig. S9B**). Further zooming in the nucleotide composition around the insertion sites did not reveal any consensus motif (**Additional file 1: Fig. S10**). ONSEN and AtCOPIA21 also generated statistically significant enrichment for euchromatin and genes in most cases, albeit not as noticeable as EVADE (**Figure 3D, E; Additional file 1: Fig. S9**).

### Epigenomic signatures define somatic integration bias of TE families

Integration preferences for some transposons have been shown to be influenced by the epigenomic state of the underlying chromatin [Quadrana et al., 2019; Cao et al., 2023; Tsukahara et al., 2025]. We therefore investigated if the sites of *de novo* somatic insertions are correlated with specific chromatin features including histone variants and modifications (source datasets provided in **Additional file 2: Table S4**). Aligning with the observed distribution confined to the chromosomal arms, somatic insertions from all the TE families showed a significant depletion in genomic regions marked by the constitutive heterochromatin mark, H3K9me2 (**Figure 4A, B; Additional file 1: Fig. S11**). *De novo* EVADE somatic insertions were enriched in genomic regions characterised by repressive facultative heterochromatin mark H3K27me3, histone variant H2A.Z that is associated with both gene activation and repression, and histone mark H3K4me1, which is a distinctive signature for active genes (**Figure 4A**). Remarkably, all other investigated active chromatin marks were significantly depleted in regions of EVADE somatic insertions (**Figure 4A**). To substantiate our findings, we assembled 2,472 genes that are commonly targeted by new EVADE somatic integrations in all applied conditions and samples, and evaluated the type and extent of chromatin signatures over their body compared to a random set of genes without EVADE insertions. We observed high signal deposition of H3K27me3, H2A.Z, and H3K4me1 over the gene bodies of EVADE-targeted genes while a contrasting trend was observed for the other active marks (**Figure 4C**).

**Figure 4.**
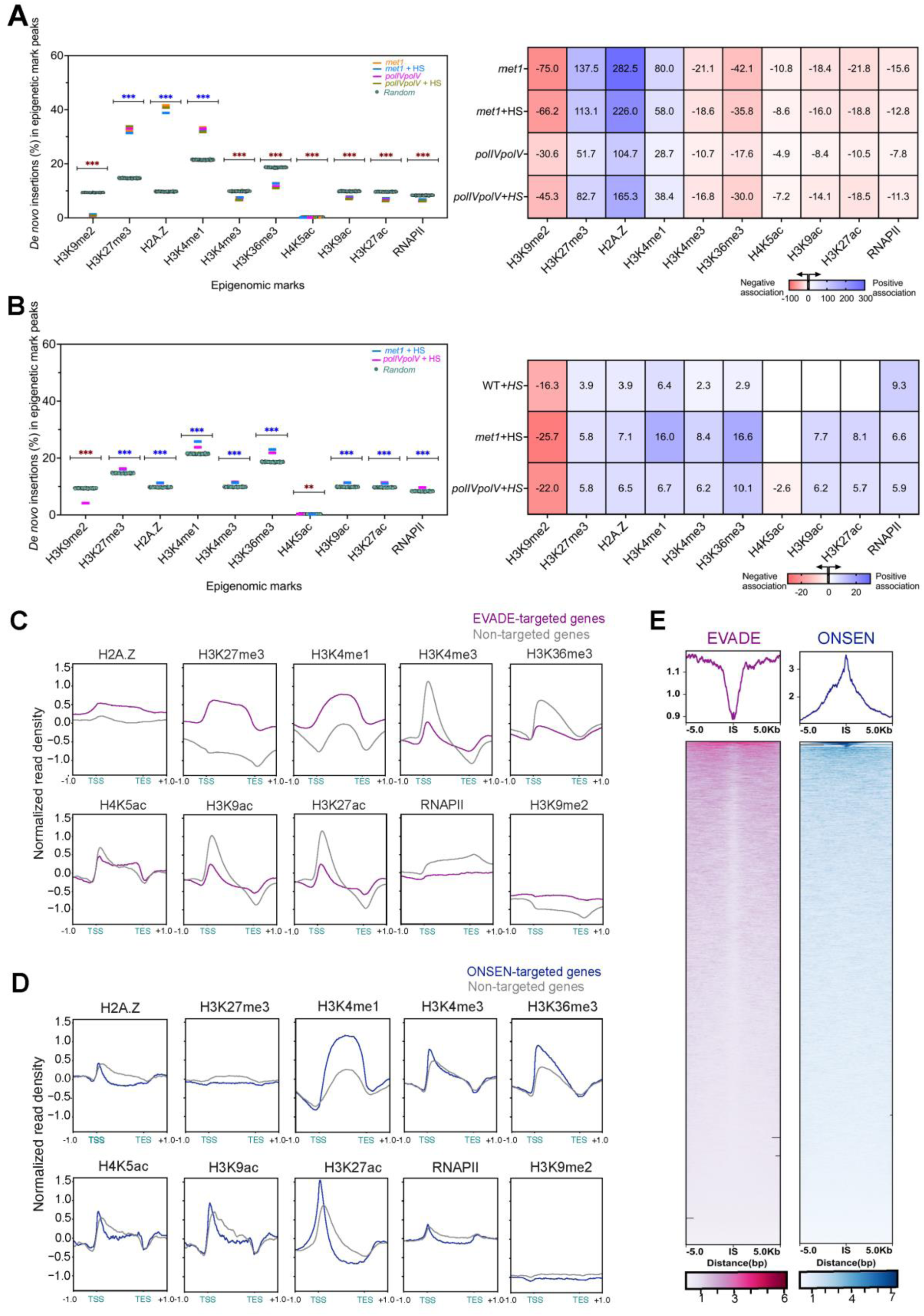
Epigenomic landscape of somatic insertion sites A, **B.** Proportional distribution of *de novo* somatic insertions of EVADE (**A**) and ONSEN (**B**) in the peak regions of various epigenomic marks (left). Colored horizontal lines represent observed values, while teal-green dots indicate expected values for the randomized insertions. Permutation test was applied to evaluate level of significance (***p< 0.001; **p<0.01). Positive association marked by blue asterisk while negative association shown by red asterisk. Heatmap depicting Z-score representing the strength of association between somatic insertions of EVADE and ONSEN, and epigenomic marks, computed through permutation tests (right). Positive association shown by blue gradient while negative association represented by red gradient. Empty blocks show samples with non-significant associations **C, D.** Metagene plots illustrating the levels of various epigenetic marks over genes targeted for *de novo* somatic insertions by EVADE (**C**) and ONSEN (**D**) compared to a randomized set of genes with no insertions from the respective TE family. **E.** Metagene and heatmap profiles at and around (± 5kb flanking region) the insertion sites of EVADE and ONSEN integration events for *met1*+HS samples generated using ATACseq data, showing an inverse tendency for the two TE families for open chromatin regions. Metagene profiles for other samples are shown in **Additional file 1: Fig. S12**. Metagene plots in C-E are generated using datasets from *met1* mutant and WT unless otherwise stated. Dataset IDs are shown in **Additional file 2: Table S4**.

Like EVADE, the somatic insertions of ONSEN showed significant integration preferences for H3K27me3, H2A.Z, and H3K4me1, yet with lower strength of association (**Figure 4B**). In contrast to EVADE, we also observed a substantial enrichment of regions occupied with RNAPII and other active chromatin marks including H3K4me3, H3K36me3, H3K27ac and H3K9ac in the mutant lines (**Figure 4B**). Notably, ONSEN integrations in wild type *Arabidopsis* did not exhibit a target preference towards two of the additional active marks, H3K27ac and H3K9ac (**Additional file 1: Fig. S11A**). The quantification of chromatin marks over 453 genes targeted by ONSEN somatic insertions across all samples by and large mirrored the genome-wide patterns. However, no obvious enrichment for H2A.Z and H3K27me3 was detected, suggesting that insertional preference of ONSEN for H2A.Z and H3K27me3 is not limited to genic targets (**Figure 4D**). To verify the contrasting integration choices of EVADE and ONSEN for active chromatin marks, we further analyzed the chromatin landscape at and around the integration sites using ATACseq data. EVADE integration sites were depleted in open chromatin regions, opposing ONSEN integration sites that were strongly associated with open accessible chromatin (**Figure 4E; Additional file 1: Fig. S12**). Finally, the integration trends of AtCOPIA21 closely resembled those of ONSEN suggesting an insertional preference for active regions. However, these associations were not significantly enriched across all active marks and samples (**Additional file 1: Fig. S11B, C**). Together, our findings provide evidence that real-time somatic transposition in *A. thaliana* presents family-specific biases towards distinct chromatin environments, expanding the spectrum of chromatin features that guide TE integration patterns.

### Insertion hotspots characterized by recurrent somatic insertions

We next investigated whether specific genomic “hotspot” regions are preferentially selected for recurrent somatic insertions. For each TE family and experimental condition, “insertion hotspots” were characterized as genomic regions (10 kb bins) with insertion frequencies 10 times higher than expected by chance. For EVADE, we detected 40 local hotspots in *met1*, 31 in *met1*+HS, 67 in *polIVpolV* and 53 in *polIVpolV+*HS (**Figure 5A; Additional data 4**). The average frequencies of the EVADE somatic integrations in the hotspot clusters ranged from 41 to 139 in *met1* and from 8 to 37 in *polIVpolV* (**Figure 5B)**. Notably, 10 genomic hotspots were persistently targeted for EVADE somatic insertions across all samples and mutant lines, suggesting stable preferential targeting of these genomic sites (**Figure 5B)**. In addition, 25 and 26 hotspots were shared between the control and heat-stressed samples of *met1* and *polIVpolV*, respectively. For ONSEN, we identified 23, 4 and 1 hotspots in heat-stressed samples of wild-type, *met1* and *polIVpolV*, which contained between 4 to 28 insertions, 18 to 56, and 19 insertions respectively (**Figure 5A; Additional data 4)**. ONSEN showed persistent targeting of a single hotspot that was shared across different samples (**Additional file 1: Fig. S13**). Notably, no hotspot loci were shared between EVADE and ONSEN. Overall, the hotspots from both TE families were uniformly distributed across all five chromosomes (**Figure 5B; Additional file 1: Fig. S13**). No hotspot clusters were identified for AtCOPIA21.

**Figure 5.**
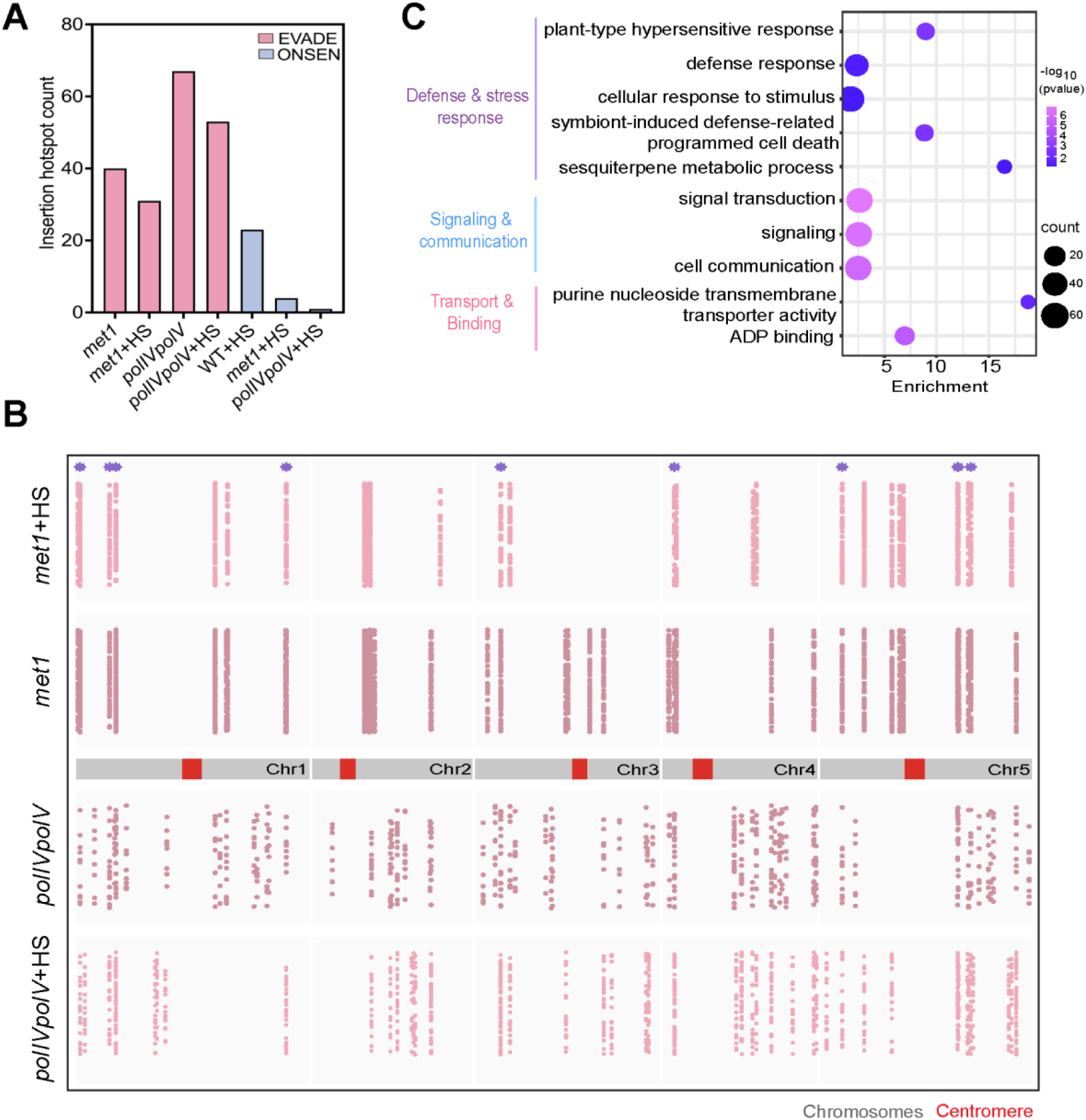
Genomic hotspots characterized by recurrent *de novo* insertions. **A.** Somatic insertion hotspots identified for EVADE and ONSEN. **B.** Distribution of insertion hotspots of EVADE across the *Arabidopsis* genome. Each dot represents an insertion in a 10kb hotspot cluster. Hotspots shared between all samples have been indicated by purple asterisk. **C.** Gene ontology analysis of genes co-localized in the EVADE hotspots. Enriched GO terms under biological function and molecular processes have been shown.

To gain insights into the biological relevance of these somatic TE hotspots, we performed gene ontology (GO) analysis on the genes co-localized within the EVADE hotspots. The principal GO terms enriched under biological processes were associated with pathways involved in defense & stress response, as well as signal & communication while for the molecular function, ADP binding and transporter activity emerged as the predominant classes (**Figure 5C**). These results mirror earlier studies showing a preferential association of EVADE germline insertions in environmentally responsive genes [Quadrana et al., 2019; Baduel et al., 2021]. No GO categories were enriched for genes within ONSEN hotspot sites.

### Assessing predisposition of environmentally-responsive regions as biased *de novo* integration sites

The specific enrichment of environmentally responsive genes within the insertion hotspots prompted us to investigate if such regions are predisposed for enhanced *de novo* TE integrations. We explored our hypothesis by analysing two functional regions in the Arabidopsis genome which are actively modulated by environmental cues: Resistance (R) genes, which mediate plant defense against pathogens [Lee & Chae, 2020] and biosynthetic gene clusters that contain multiple genes with roles in specialized metabolic pathways and show co-regulation upon environmental and stress cues [Nützmann et al., 2016]. The insertion density of *de novo* somatic transposition overlapping with these regions was analyzed and compared to a similar set of randomly generated loci, sampled 1000 times by applying permutation tests.

Focusing on 170 R genes in *A. thaliana* [Lee & Chae, 2020; Yang et al., 2020], we detected high rates for somatic insertions of both EVADE and ONSEN (p<0.01, Permutation test; **Figure 6A; Additional data 5**). The SSI4 and RPS4 gene clusters exhibited the highest frequencies of EVADE somatic insertions, with a total of 370 and 199 independent integrations across all samples (**Figure 6B; Additional data 5**). Reflecting patterns observed for R genes, somatic insertions for EVADE - but not ONSEN - were substantially over-represented in the 62 biosynthetic genes in *met1* and heat-stressed *polIVpolV* (p<0.01, Permutation test; **Figure 6C, D; Additional data 6**). In fact, the epigenomic profile of both the RPS4 and thalianol clusters showed strong enrichment for the same marks (H3K27me3, H2A.Z, and H3K4me1) that were associated with the general EVADE insertion bias (**Figure 6B, D)**.

**Figure 6.**
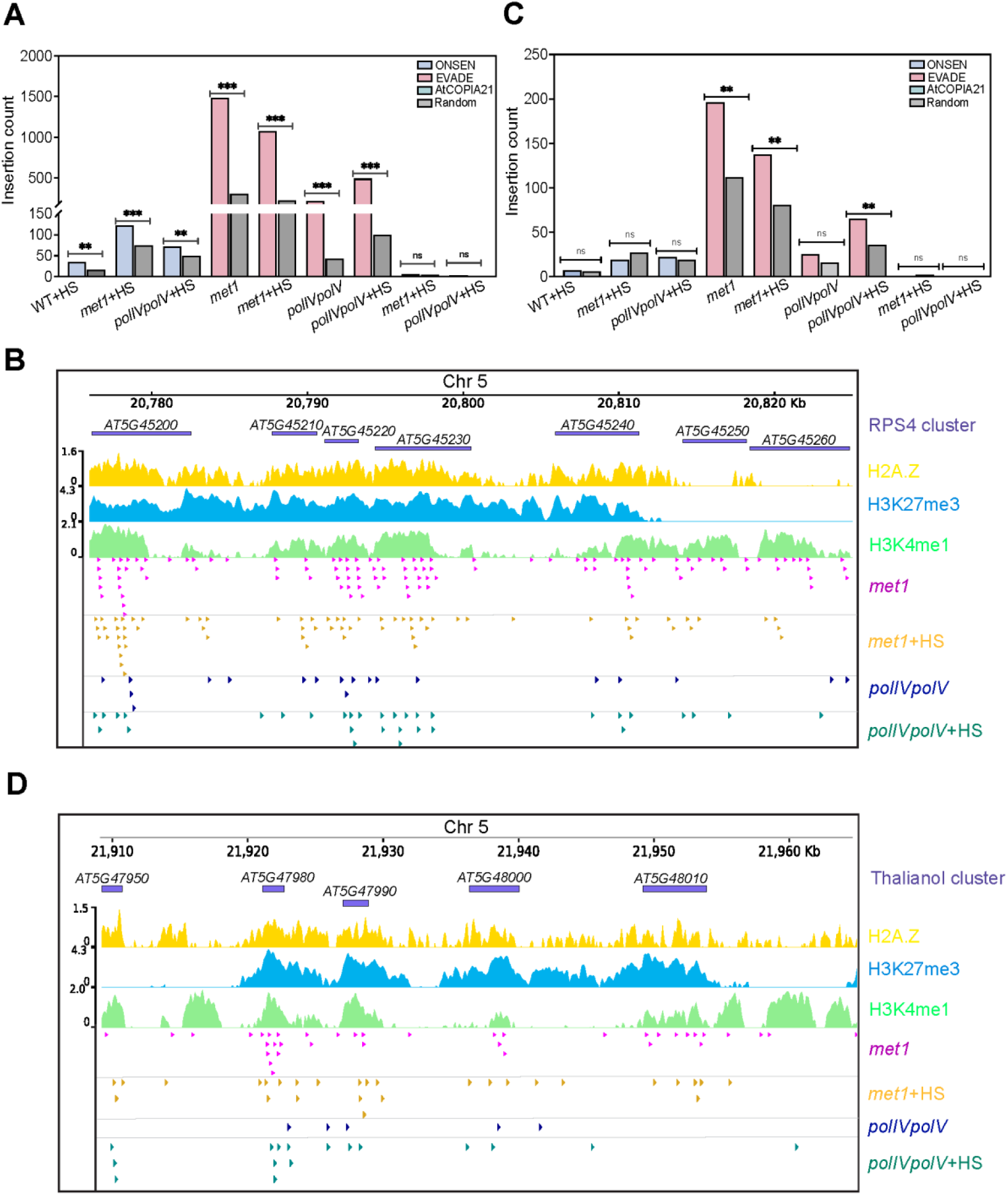
Environmentally-responsive regions as biased *de novo* integration sites. A, **C.** *De novo* somatic insertions of the three TE families overlapping with R genes (**A**) and biosynthetic genes (**C**). Permutation test was applied to evaluate level of significance (***p<0.001; **p<0.01; ns-non-significant). **B, D.** Genome browser profiles showing distribution of *de novo* somatic insertions of EVADE in representative R gene cluster, RPS4 (B) and thalianol biosynthetic gene cluster (D) of *A thaliana*. Tracks showing the distribution of genes (purple) and epigenomic marks including H2A.Z (yellow), H3K27me3 (blue) and H3K4me1 (green) are presented. Each arrowhead in the four tracks below indicates a unique somatic insertion event of EVADE identified in samples from mutant lines. Genomic coordinates have been indicated for each plot.

Clusters of both R genes and biosynthetic genes have previously been reported to be enriched for TEs [Boutanaev and Osbourn, 2018; Kawakatsu et al., 2016; Teasdale et al., 2024]. To test if other regions of high TE load are preferentially targeted by *de novo* TE insertions, we examined and contrasted the distribution of somatic insertions within the KNOT region - a 3D nuclear structure that has been demarcated as a preferred integration hotspot for invading TEs [Grob et al., 2014; Sun et al., 2020]. However, we found no enrichment for new somatic insertions of all three TE families in the KNOT regions (**Additional file 1: Fig. S14**; Permutation test, p>0.05). Our findings show that resistance and biosynthetic genes function as preferential integration sites for TEs, therefore suggesting that TEs may have a critical role in shaping their evolution and plastic functionality.

## Discussion

In multicellular organisms, somatic cells do not share identical genomes, instead they exhibit genomic mosaicism with significant differences between cells [Schoen and Schultz, 2019]. The genome-wide contribution and impact of TEs in this mosaicism has been mostly studied in animals, focusing on brain cells and cancerous or ageing tissues [Baillie et al., 2011; Gorbunova et al., 2021; Siudeja et al., 2021; Yang et al., 2022; Bodea et al., 2024]. Here, our study provides direct evidence for TE-mediated genomic variation in somatic cells of a plant at an unprecedented scale. By identifying hundreds of thousands of *de novo* somatic transposition events (**Figure 2**), we show that the transcriptional activity of TEs rapidly translates to integration of TE copies into the genomes of somatic cells, thus generating genetic diversity between cells. Although transient, the variability contributed by somatic TE insertions could influence intra-organismal cell identity and tissue functionality, in addition to imparting detrimental or useful effects. For example, evidence that somatic TE insertions could confer beneficial phenotypic variation in plants is reflected by the vast diversity observed in the bud sport somatic chimeras of clonally propagated crop species such as apple and oranges [Foster & Aranzana, 2018; Wang et al., 2021b; Ban et al., 2022; Wu et al., 2024]. Given that our sampling consisted only of a few leaves from young seedlings, we speculate that somatic transposition in the plant body is an explosive and potentially recurring event when conditions are favorable in both natural populations and experimental conditions.

Recent work illustrates how stress-activated ONSEN elements can modulate the adaptive response of genes near a novel fixed insertion site [Roquis et al. 2021; Raingeval et al., 2024]. As we provide evidence that ONSEN and other TEs actively transpose to new genomic sites in somatic cells of Arabidopsis including wild-type plants, it is likely that *de novo* TE insertions also imitate this modulated stress response of proximal genes in somatic cells and thereby, transiently alter stress physiology in a spatio-temporally dynamic manner. The downstream effects of this modulation can be important for both crops and perennial species like trees, and also in species with high loads of recently active TEs that can collectively contribute en masse to somatic mosaicism. Very recently, one study in maize attempted to quantify levels of somatic transposition of the Mutator DNA transposon using ultra-deep short-read sequencing but with no analysis of their genome-wide genetic and epigenetic characteristics [Scherer et al., 2025], while two studies in *A. thaliana* retrieved a small number of somatic insertions using different long-read sequencing approaches [Movilli et al., 2025; Merkulov et al., 2023]. Along with our findings, these studies underscore an emerging focus on investigating the dynamics of TEs in the soma of plants, warranting a systematic investigation across diverse plant systems.

Our work demonstrates that global somatic transposition rates as well as integration sites are TE-specific and influenced by the genetic background of the host. In line with previous reports highlighting EVADE as the most successful TE in generating a high frequency of germline insertions in Arabidopsis [Quadrana et al., 2016; Quadrana et al., 2019], we observed extreme mobility of EVADE in somatic cells of epigenetic mutant lines (**Figure 2**). For ONSEN, abundant new somatic insertions were recorded in heat-stressed samples of both wild-type and mutant Arabidopsis lines. Compared to EVADE and ONSEN, the somatic transposition rate of AtCOPIA21 was low in the studied mutants, which is consistent with TE families responding with varying levels to induction triggers [Liu et al., 2022]. For all families, loss of CG methylation in *met1* imparts greater release of repression than in the double *polIVpolV* mutant. This disparity likely stems from the central role of *met1* in maintaining TE suppression, compared to the locus-specific and reinforcing action of *polIVpolV* and RdDM [Panda et al., 2016]. Furthermore, loss of *met1* not only attenuates DNA methylation over CG sites but also modifies several chromatin signatures necessary for maintenance of genome-wide TE repression, which could explain the pronounced activation of TEs observed in this mutant [Reinders et al., 2009].

Delimiting the factors that guide TEs to their initial integration sites is fundamental to TE biology but also crucial for our broader understanding of genome dynamics. In the absence of selection during somatic transposition, we found that all three TE families showed a general *de novo* integration preference for the euchromatin-rich chromosomal arms and depletion from constitutive heterochromatic regions in the peri/centromere (**Figure 3**) - a pattern that aligns with previous observations for non-reference segregating insertions for these TE families [Gaubert et al., 2017; Quadrana et al., 2019; Roquis et al., 2021]. At the regional scale, we observed deviating integration site preferences with EVADE showing much higher integration preference for genes compared to ONSEN and AtCOPIA21. At the epigenetic level, previous reports have demonstrated the critical role of H2A.Z and H3K27me3 chromatin marks in shaping the preferred integration niche for germline insertions of Copia families [Quadrana et al., 2019; Roquis et al., 2021]. Here, we also observed that *de novo* somatic insertions of EVADE and ONSEN were enriched over H2A.Z and H3K27me3 marked regions, albeit with notably stronger enrichment for EVADE (**Figure 4**). In addition, both families have a strong integration preference for H3K4me1, which is a conserved chromatin mark enriched over active genes and associated with transcription [Oya et al., 2022]. H3K4me1 is also actively involved in the recruitment of the DNA repair machinery and has been correlated with lower mutation rates in Arabidopsis [Monroe et al., 2022; Quiroz et al., 2024], which raises the possibility that H3K4me1-labelled sites are inherently susceptible to new TE integrations and, therefore, are targeted by the DNA repair machinery to facilitate removal of new integrations and protect genome integrity.

In contrast to ONSEN, EVADE somatic insertions were depleted from active chromatin marks other than H3K4me1 and from areas of open chromatin. This finding is consistent with TEs occupying different target site niches at the local epigenetic level, helping to avoid conflict for space in the genome ecosystem. This spacing may increase mutational load, depending on epigenetic state and environmental condition. Indeed, the EVADE and ONSEN hotspots showed no overlap, yet some genomes areas were highly susceptible to insertions of both families (**Figure 6**). For example, within R gene clusters, we observed hundreds of recurrent and independent insertions of EVADE and ONSEN TEs. This may suggest that R genes are in fact acting as sponges for new somatic TE insertions, which may drive the diversification of these disease resistance loci, and may modulate their regulatory properties. Intriguingly, EVADE expression is elevated upon pathogen stimulation [Yu et al., 2013; Zervudacki et al., 2018], and so may directly affect R gene cluster architecture and expression in individual cells and tissues [Zervudacki et al., 2018; Lai et al., 2020]. Our findings demonstrate that R gene clusters are predisposed for TE integrations and that this feature may be a key element in the adaptability of defence genes to changing pathogen threats.

In summary, by documenting the accumulation en masse of TE insertions in the soma of Arabidopsis, we provide evidence for the plasticity of plant genomes at the cellular level but also for the unique integration site preferences of TEs independent of selection. We speculate that these features enable host genomes to generate genetic diversity at genomic sites of need, which leads to genetic cellular plasticity at times of TE mobilisation. Our work establishes that studying somatic transposition is a reliable and powerful approach towards understanding TE dynamics, and, thereby, genome function and evolution.

## Methods

### Plant materials and experimental growth conditions

*A. thaliana* wild type Col-0 and mutant plants were grown on ½ Murashige and Skoog (MS) medium in a growth chamber at 22°C under long day conditions (16h light, 8h dark) following an initial stratification for 2 days at 4°C. Details of the mutant alleles used in the study are provided in the **Additional file 2: Table S1**. All the wild type and mutant alleles were in Col-0 ecotype background. Young leaves harvested from a pool of ∼100 individual seedlings were used for all the experiments.

### Heat stress regimes for AtCOPIA78 (ONSEN) induction

For heat stress application, the 10 days old seedlings grown on MS medium were subjected to three variable heat regimes to quantify ONSEN induction (**Figure 1A, Additional file 1: Fig. S1**). A high temperature of 37°C was used for all heat-stress treatments.

i. For the present study, we define long-duration heat stress as high temperature exposure that spanned over 8 continuous days. The WT seedlings pre-treated to low temperature (6°C) were allowed to grow for 8 days at high temperature. At every 24 hrs, leaf samples were harvested to monitor ONSEN activity. Additionally, at each indicated time point, a similar set of HS seedlings were transferred to normal growth conditions (22°C, 16h light, 8h dark cycle) for a 48hrs recovery period (**Figure 1A**).
ii. Effect of low temperature before heat application on ONSEN activation was tested by exposing the WT seedlings to high temperature of 37°C for 48 hrs with or without prior exposure to chilling temperature of 6°C for 24 hrs (**Additional file 1: Fig. S1**).
iii. In the third variation of the applied heat regime, cold-treated WT and mutant seedings were exposed to high temperatures for up to 48 hrs. Samples were collected and analyzed at intervals of 0, 6, 12, 24 and 48 hrs during the heat stress period (**Figure 1A**).

Leaf tissues were harvested immediately after completion of specified timepoints for each treatment and frozen in liquid nitrogen. The samples were stored at-80°C until further use.

### RNA extraction and qRT-PCR for TEs transcription abundance

Total RNA was extracted from harvested tissues using Trizol reagent (Sigma) and DNase-treated with Turbo DNA-free kit (Thermo) following manufacturer’s instruction. Starting with 1µg of total RNA, complementary DNA (cDNA) was synthesized with iScript cDNA synthesis kit (Biorad) according to manufacturer’s instructions. TE transcript abundance was evaluated through qRT-PCR using PowerUp SYBR Green Master Mix (Applied Biosystems). TE primer sequences are listed in **Additional file 2: Table S5**. Expression levels were determined relative to the housekeeping control gene (18S rna or ACT2), applying the 2–ΔΔ*Ct* method. Mean and standard deviations were inferred from three biological replicates for all experiments unless stated otherwise.

### DNA extraction

For eccDNA assays and whole genome sequencing, total DNA was extracted from sampled tissues using standard CTAB method [Murray & Thompson, 1980]. For TEDseq, high molecular weight DNA was prepared after nuclei isolation from harvested tissue. Briefly, 0.5gm of leaf tissue was homogenized in a mortar and pestle using liquid nitrogen. ∼20ml of chilled nuclei isolation buffer (NIB: 20mM Tris-HCl pH 7.8, 250mM sucrose, 5mM MgCl2, 5mM KCl, 40% Glycerol, 0.25% Triton X-100, and 0.1% β-mercaptoethanol) was added to the grounded powder. The thawed homogenate was filtered through four layers of Mira cloth and a 40-μm-thick nylon mesh and the filterate was centrifuged at 6000xg for 15 mins at 4 °C. The pellet was gently washed with 20 ml of NIB and the samples were incubated on ice for 10 mins followed by centrifugation at 6000xg for 15 mins at 4 °C. This step was repeated 4 times or until a clear pellet was obtained. Next, the nuclei pellet was resuspended in prewarmed Carlson lysis buffer (100mM Tris pH 8, 2% CTAB, 1.4M NaCl, 1% PEG, 20mM EDTA) along with 25μl of β-mercaptoethanol and 40μl of RNaseA. The samples were incubated at 65°C for 1 hr with intermittent mixing. Post incubation, the contents were allowed to reach room temperature, an equal volume of chloroform:isoamyl alcohol mix (24:1) was thoroughly mixed with it and centrifuged at 6000xg for 10 mins at 4°C. The upper aqueous layer was extracted in a fresh tube and another round of chloroform:isoamyl alcohol treatment was performed. DNA was precipitated with one-tenth volume of 3M sodium acetate (pH 5.2) and chilled isopropanol. The DNA pellet was washed with 70% ethanol, air dried, and dissolved in molecular biology grade water. DNA quality was assessed through electrophoresis on 1% agarose gel and Nanodrop 2000 (Thermo Scientific) measurements. DNA amount was quantified by dsDNA High Sensitivity Assay kit and Qubit Fluorometer (Invitrogen).

### Detection of extra-chromosomal circular DNA (eccDNA) and inverse PCR

To selectively remove linear DNA and enrich eccDNA, 1µg of RNased DNA was digested with 1µl of Plasmid-Safe ATP-dependent DNase (Lucigen) in a total reaction volume of 50µl and incubated at 37°C for 16 hrs. Complete removal of linear DNA was ensured by adding an additional 1µl of DNase and ATP to the reaction mixture after 16 hrs. This step was repeated twice at 2-hour interval resulting in a total digestion duration of 20 hrs after which the DNase activity was terminated by incubation at 70°C for 30 mins. eccDNA was retrieved by overnight precipitation with 0.1 volume of 3M sodium acetate (pH 5.2), 2.5 volumes of pure ethanol and 1µl of glycogen (Thermo) at-20°C. The precipitate was centrifuged at 14,462xg for 1 hr at 4°C followed by pellet washing with 70% ethanol. The air dried eccDNA pellet was resuspended in TempliPhi Sample buffer and rolling circle amplification was performed using TempliPhi Amplification kit (Cytiva) by incubation at 28°C for 65 hrs. The reaction was terminated by inactivating enzyme at 70°C for 30 mins. The enriched eccDNA product was used as template for inverse PCR to selectively amplify TE-specific products using outward primers (**Additional file 2: Table S5**). The eccDNA circle (Chr2: 3416272-3417604_+; hereafter designated as eccUni) identified ubiquitously in multiple Arabidopsis tissues was used as a positive control for eccDNA profiling [Wang et al., 2021a]. Inverse PCR products of AtCOPIA78 (ONSEN) and AtCOPIA93 (EVADE) were cloned into pGEMT vector (Promega) and sequence was verified through Sanger sequencing.

### Whole Genome sequencing with Illumina and detection of *de novo* transposon insertion from resequencing data

HMW genomic DNA was used for preparation of Illumina short read sequencing libraries with an insert size of 350bp using TruSeq DNA Library Prep kit. The libraries were sequenced on NovaSeq6000 platform in a paired end (PE) mode (2 × 150bp) at Novogene, UK to achieve an expected genome coverage depth of over 100 folds. The ColCEN, telomere-to-telomere assembly of Arabidopsis Col-0 accession and its associated annotations [Naish et al., 2021] were used consistently throughout the study for all genomic-based analysis. The quality assessment of the raw reads was performed with FASTQC [https://github.com/s-andrews/FastQC]. Trimmomatic [Bolger et al., 2014] was used to filter the low-quality and trim adapter-contaminated reads applying the following parameters ILLUMINACLIP:TruSeq3:2:30:10 LEADING:20 TRAILING:20 SLIDINGWINDOW:4:20 and MINLEN:36. The consensus TE sequence library for ColCEN was derived by lifting over the TE annotation from Col-CC reference assembly (https://github.com/oushujun/TAIR12-TE/tree/main/data/gff) using liftoff version1.6.3 [Shumate and Salzberg, 2021]. The retrieved TE library was used as an input for TEMP2 v 0.1.6 [Yu et al., 2021] to detect *de novo* somatic TE insertions through module insertion at default settings. TEMP2 output classify the obtained non-reference insertions into three main classes: *1p1 insertions* have read support at both ends of a TE, 2p insertions have 2 or more read support only at one end of TE while singleton insertions are supported by a single read. 1p1 and 2p insertions are regarded as fixed heritable insertions while singletons as *de novo* somatic events. We re-classified the obtained *de novo* somatic insertions based on the following criteria: The insertions (classified as “1p1” and “2p” in TEMP2 pipeline) where the coverage was less than 5 reads were classified as *de novo* somatic insertions along with those supported by singletons. Further details of WGS libraries are provided in **Additional file 2: Table S6.**

### Transposable element display sequencing (TEd-seq)

TE-specific enrichment libraries were prepared by optimizing the TEd-seq protocol [Vendrell-Mir et al., 2025] for three LTR-TE families: ONSEN, EVADE and AtCOPIA21 using NEBNext^®^ Ultra™ II DNA Library Prep Kit for Illumina^®^ (NEB). Briefly, HMW genomic DNA was extracted from leaves of a pool of ∼100 Arabidopsis seedlings. To obtain fragmented DNA in a size range of 300-700bp, ∼2µg of genomic DNA dispensed in 100µl TE buffer was sonicated for 8 cycles of 30sec treatment/ 30sec delay in Covaris M220 (Peak power 75W, Duty factor 10 and 200 cycles per burst). End repair and A tailing of 1µg sheared DNA was performed using NEBNext Ultra II End Prep Enzyme Mix in a total volume of 60µl (NEB). TEd-seq custom adapters (**Additional file 2: Table S7**) were ligated using NEBNext Ultra II ligation Master Mix and NEBNext Ultra II ligation enhancer followed by size selection with AMPure XP Beads (Beckman Coulter). First round of selective TE-fragment enrichment PCR (primary PCR) was performed with outer TE-specific primer (TE*_rev; * indicate TE family; **Additional file 2: Table S7)** and adapter-specific primer in a total volume of 50µl with the following cycling conditions: 98°C for 30 sec followed by 20 cycles of 98°C for 10 sec, 61°C for 75 sec and a final cycle at 61°C for 5 min. The PCR products were cleaned and size-selected using 0.9X ratio of AMPure XP Beads. The cleaned PCR product was used as a template for a second round of nested PCR using an inner TE primers mix with Illumina P5 overhang (P5_TE*; * indicate TE family) and P7 adapter primer with index in a total volume of 25µl. The amplification conditions used are: 98 °C for 30 sec, *n* cycles of 98°C for 10 sec, 61°C for 75 sec, *m* cycles of 98°C for 10 sec, 72°C for 75 sec and final cycle at 72°C for 5 mins (*n* and *m* represents specific PCR cycle number optimized for each TE family; **Additional file 2: Table S7**). The final DNA library was subjected to two-sided size selection to capture fragments in range of 200-600bp with AMPure XP beads using sequential ratios of 0.67X and 1X, and eluted in a final volume of 30µl TrisHCl (10mM). Illumina short-read sequencing was performed at Novogene,UK on NovaSeq platform in a paired end (PE) mode (2 × 150bp). For pooled multiplexed TEd-seq libraries, primers of different targeted TEs were mixed in equimolar concentrations. Library details are given in **Additional file 2: Table S3**.

### Computational analysis with deNOVOEnrich

The TEd-seq bioinformatic pipeline as described in the original reference [Vendrell-Mir et al., 2025] is intended to capture non-reference segregating insertions. To efficiently identify and profile rare somatic transposition events in addition to non-reference segregating TE insertions from TEd-seq data, we have designed a new computational pipeline, deNOVOEnrich [https://github.com/hAmbreen02/deNOVOEnrich]. The general principles and overall workflow of the deNOVOEnrich pipeline is presented in **Additional file 1: Fig. S5**.

For deriving transposition rate across different samples, we normalized the raw data for each TE family by downsampling reads in the samples to the scale of the sample with least number of TE-specific reads. The first step of *deNOVOEnrich* pipeline involves stringent identification of TE-specific paired-end reads from TEd-seq data. This is achieved by using terminal end sequence (∼50-60 bp) of the targeted TE as a bait, which sensitively detect TE-associated reads while promoting filtering of non-specific amplification or chimeric reads from the analysis. Additionally, this step mediated through sequential deployment of two filtering algorithms, cutadapt [Martin, 2011] and fastp [Chen S., 2023] also identifies low quality reads, sequencing artefacts (including reads with polyG tails associated with NovaSeq technologies) and short read fragments (<20 nt in length). The identified TE-specific reads are then retrieved from the original raw data with seqtk [https://github.com/lh3/seqtk] and used for further processing after adapter trimming with trimmomatic with the parameters: ILLUMINACLIP:TruSeq3:2:30:10 LEADING:3 TRAILING:3 SLIDINGWINDOW:4:15 and MINLEN:20 (Bolger et al., 2014). To further increase alignment accuracy and detection of TE-genome split junction, we generated longer DNA contigs by concatenating read pairs into single reads with a minimum overlap of 5 base pairs at their ends and keeping the contig lengths in a range between 50 bp (-n) to 300 bp (-m) using PEAR (Zhang et al., 2014). The long DNA contigs are then mapped onto the reference genome with bowtie2 (Langmead et al, 2012) allowing split-read mapping in local mode (--local, –very-sensitive). Subsequently, alignments on the reference genome are filtered to retain only those mapped reads that align at a unique location in the genome and harbored at least 40 nucleotides of soft-clipped sequence while rejected reads that mapped on multiple locations and/or lacked sufficient split support. PCR duplicates were marked with picard MarkDuplicates (http://broadinstitute.github.io/picard/). Simultaneously, the merged DNA contigs were also mapped separately on TE extremity sequence (enclosing TE region from primer site to 5’ extremity end) in local mode with bowtie2. Next, we preserved split contigs by establishing shared read homology between the target TE and reference genomic sites using Readtagger with default parameters [Siudeja et al., 2021]. Using the SAM tags assigned by the Readtagger, TE-specific tagged reads that mapped to both the reference genome and targeted TE sequences were filtered. These TE-tagged alignments which are indicative of TE:genome break sites were then used to determine *bona-fide* non-reference TE insertion sites in the genome.

TE insertions were mainly classified into two categories: *de novo* somatic insertions and non-reference segregating insertions, depending on the read depth at the putative TE integration sites. Since somatic insertions occur in low frequency in single or few cells, which depends on the number of cell replication cycles during our experimental conditions, all insertions supported by a low total read depth of ≤5 including singleton reads were classified as *de novo* somatic insertions. TE insertions with a threshold coverage of ≥30 reads, supported by at least 5 unique contig reads (sharing the same start site but variable stop sites) and with at least 3 of the unique reads containing 3 or more PCR duplicates were classified as non-reference segregating insertions.

Highly stringent filters were applied at each step of the pipeline to ensure removal of false positives and reduction of background noise signal. Firstly, TE insertions localized within 1kb window of reference TE sites in the genome were discarded as previous studies have shown that these regions contribute to high rate of chimeric read generation leading to frequent misalignment [Treiber & Waddell, 2017]. Next, insertions within 50bp window of mono-nucleotide genomic stretch or overlapping with organelle genomes were filtered to remove ambiguous alignment sites. Finally, for *de novo* somatic insertions, we discarded insertion sites commonly recorded between different samples as the likelihood of shared somatic insertions is highly improbable [Zhao et al., 2019]. The source code of the pipeline is available at the GitHub repository [https://github.com/hAmbreen02/deNOVOEnrich]. Genomic distribution of *de novo* TE insertions was displayed through circular plot generated through Circos [Krzywinski et al., 2009].

### Genomic and epigenomic features association for TE-targeted sites

Target site preferences of different TE families were studied by evaluating the overlap between TE insertions and several genomic feature datasets. To analyze integration preferences at the genomic regional level, the Arabidopsis genome was differentiated into following distinct regions: protein coding genes (PCGs), proximity to genes (500 basepairs upstream or downstream of PCGs), reference transposable elements and intergenic regions. The frequency proportion of *de novo* integrations overlapping in each genomic region was recorded. Further, insertions in the PCGs were classified into different genic categories in priority order of 5’UTR, 3’UTR, exons and introns. To determine the significance levels of insertion bias, permutation statistical test was applied for overlaps between somatic insertions and genomic compartments using randomizeRegions function in regioneR [Gel et al., 2016]. The degree of enrichment is indicated by permutation Z-scores. Analysis of sequence motifs at the integration sites was performed by retrieving five nucleotides flanking both the upstream and downstream region of the insertion sites with bedtools utilities, *slop* and *getfatsa* [Quinlan & Hall, 2010]. The sequences were used as input for motif search using meme [Bailey et al., 2015]. To determine local sequence patterns around the insertion sites, sequence logo was generated with WebLogo3 [Crooks et al 2004]. Epigenomic datasets from ChIP-seq and ATAC-seq (**Additional file 2: Table S4**) were processed by first treating the raw reads for removal of low quality and adapter contaminated reads with Trim Galore with options--length 30--quality 20--stringency 1-e 0.1 [https://github.com/FelixKrueger/TrimGalore]. The high-quality reads were aligned to ColCEN reference genome with bowtie2 using parameters:--end-to-end--sensitive. PCR duplicates were removed by samtools markdup. For ChIP-seq, peak calling was performed using macs3 with the following options-q 0.00001,-f BAMPE [Zhang et al., 2008]. Depending on the epigenetic mark, either broad or narrow peak calls were used for evaluating its association with *de novo* insertions based on overlapping frequency. For ATACseq, findPeaks function of HOMER was used for calling peaks applying following parameters:-region-size 200-minDist 202-gsize 135000000-tbp 0 (http://homer.ucsd.edu/homer/). Normalization and visualization of ATAC profiles was performed using deepTools [Ramirez et al., 2014]. Metagene plots of epigenomic features were generated using *computeMatrix*, *plotProfile* and *plotHeatmap* utilities of deepTools.

### Insertion hotspots

TE insertion hotspots were defined following Wang et al., 2018. Briefly, identification of insertion hotspots was performed by dividing the Arabidopsis reference genome into equal-sized bins of 10kb with bedtools makewindows followed by evaluation of insertion coverage per bin with bedtools coverage. Based on the hypothesis of no integration bias, the expected number of insertions per 10 kb window were calculated. Depending on the transposition rates, bins harboring 10 folds higher insertions than that is expected by chance were classified as insertion hotspots. Gene ontology enrichment analysis was performed for genes localized in insertion hotspots using PANTHER Classification System (Mi et al., 2013).

### Analysis of TE-populated genomic regions: R genes, biosynthetic genes and KNOT region

The Arabidopsis functional genomic regions including R genes [Lee & Chae, 2020; Yang et al., 2020], biosynthetic genes [Yu et al., 2016] and KNOT-regions [Grob et al., 2014] were assessed for their pre-disposition efficiencies as potential sites for attracting new TE integrations. To assess TE integrations bias in these regions, loci were randomized while preserving their intrinsic structure and characteristics, and association analysis was performed based on permutation tests with regioneR [Gel et al., 2016]. For R genes and biosynthetic genes, all protein coding genes were provided as universe for resampleRegions functions. Genomic insertion distribution plots were generated with pyGenomeTracks [Lopez-Delisle et al., 2021].

### PCR validation of non-reference segregating insertions

For validation of the genomic sites of non-reference segregating insertions (**Additional file 1: Supplementary Information 1**), we randomly selected four TE integration sites (FTEins_1-4). Primers were designed from the 3’ end of the TE sequence and the putative flanking genomic site to capture the 3’ break junction of the TE insertion (**Additional file 2: Table S5**). PCR was performed on 0.5ng of DNA pooled from 100 seedlings with following amplification conditions: 96 °C for 3 min, 25 cycles of 96°C for 30 sec, 60°C for 30 sec, 72°C for 60 sec and final cycle at 72°C for 5 mins.

## Declarations

### Ethics approval and consent to participate

Not applicable

### Consent for publication

Not applicable

### Availability of data and materials

All sequencing data generated in the present work including whole genome sequencing and transposon display sequencing (TEd-seq) have been deposited in the NCBI Sequence Read Archive (SRA) (https://www.ncbi.nlm.nih.gov/) under the BioProject number PRJNA1270934. The source code for deNOVOEnrich and additional files employed for the analysis are available on Github: https://github.com/hAmbreen02/deNOVOEnrich. Details of the public databases used for the epigenomic analysis are provided in Additional file 2: Table S4.

### Authors’ contributions

H.A., A.B. and H.W.N conceived and designed the study. H.A. performed all experiments and computational analysis. R.K.S. contributed the plant material. B.L. and L.Q. supported TEd-seq experiments and data analysis. H.A., A.B. and H.W.N wrote the manuscript with input from R.K.S. and L.Q. All authors read and approved the manuscript.

## Acknowledgements

We are grateful to Pierre Baduel and Pol Vendrell for valuable discussion on the experimental setup, Jessica Taylor for her help in cataloguing the Resistance (R) genes, the technical team at the School of Life Sciences, University of Sussex for their support in setting up the laboratory work and members of the Nützmann lab for discussion and suggestions.

## Supplementary Information

**Additional File 1:** Supplementary information 1 and Supplementary Figures S1-S15

**Additional File 2**: Supplementary Tables S1-S7

**Additional data 3:** Somatic insertions from different TE families recorded through whole genome sequencing

**Additional data 4:** Somatic insertion hotspots identified for EVADE and ONSEN.

**Additional data 5:** List of 170 R genes and number of somatic insertions observed in each gene

**Additional data 6**: List of 62 biosynthetic genes and number of somatic insertions observed in each gene

