## Additional File 1 for "Somatic mobility of transposons is explosive and shaped by distinct integration biases in *Arabidopsis thaliana*"

**Supplementary Information 1**

**Non-reference segregating insertions in Arabidopsis mutants**

In addition to *de novo* somatic insertions that are generated and detected within a single generation, our pipeline also captured non-reference segregating insertions based on the high read coverage of the TE-genome junctions (≥30 reads, see Methods) (**Figure 2A,B**). In our experimental setup, these insertions could have only occurred in the two rounds of seed propagation prior to the experiments. For ONSEN, the seedlings or their parental lines were never subjected to heat-stress up to the point of the TEd-seq library preparation, so no insertions were identified, which matches our expectations. In addition, no insertions were observed in wild-type plants for EVADE and AtCOPIA21, underscoring the lack of spontaneous TE mobilization and also the low FDR in our approach. We identified, however, several non-reference segregating insertions for EVADE, especially in *met1* with (185) and without (284) heat-stress (**Fig. S15A**). Fewer insertions were observed in *polIVpolV* (164 and 105 respectively) (**Fig. S15A**), which is consistent with an overall lower transposition capacity of EVADE in *polIVpolV* compared to *met1* (**Figure 2E**). We captured AtCOPIA21 segregating transpositions only in *met1*, with 22 (no-stress) and 8 (heat-stress) detected insertions (**Fig. S15A**). We validated four segregating insertions with primers designed from the TE end and the flanking sequence in non-stressed *met1* plants (**Fig. S15B,C**).

**Supplementary Figures**

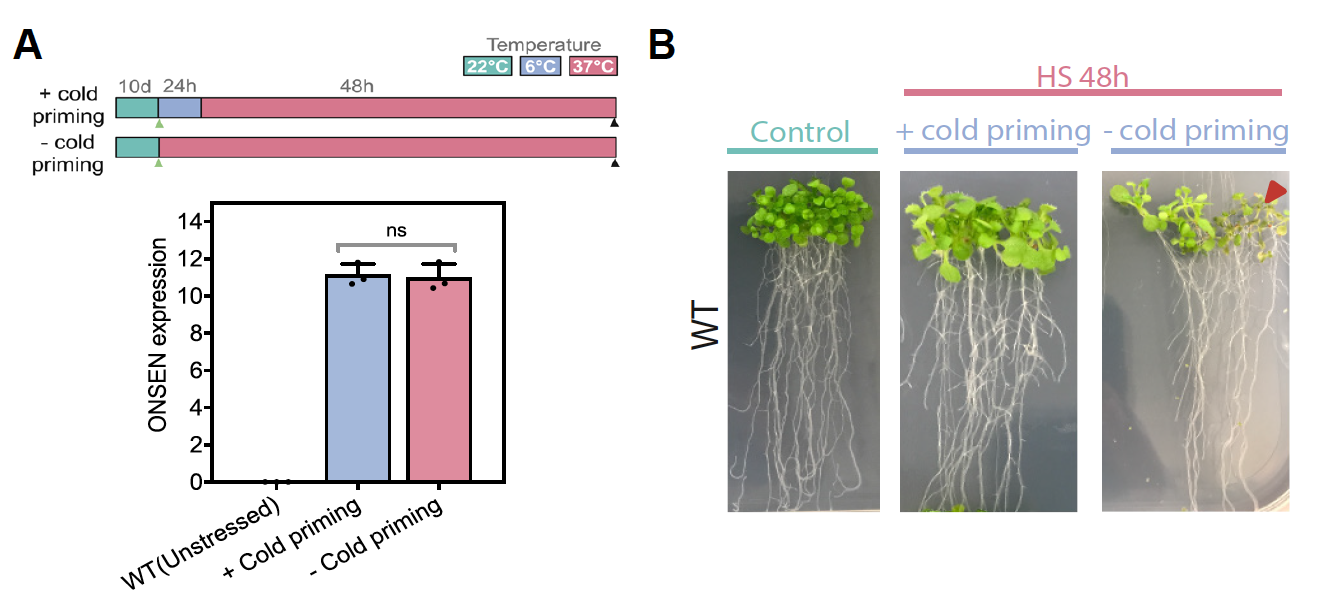

**Fig. S1. A.** Schematic illustrating experimental conditions adopted for evaluating the effect of cold priming on ONSEN transcription upon subsequent heat stress induction. Sampling points have been indicated by green (control) and black (heat-stressed) arrowheads in the schematic. Relative expression level (mean ± s.d. plotted) of ONSEN in wild-type Arabidopsis (Col-0) seedlings with and without cold priming (n=3). *Student* *t-test* applied to evaluate level of significance (ns: non-significant). **B.** Plant phenotype studied for effect of cold priming prior to HS. Necrotic plant tissue indicated by red arrowhead.

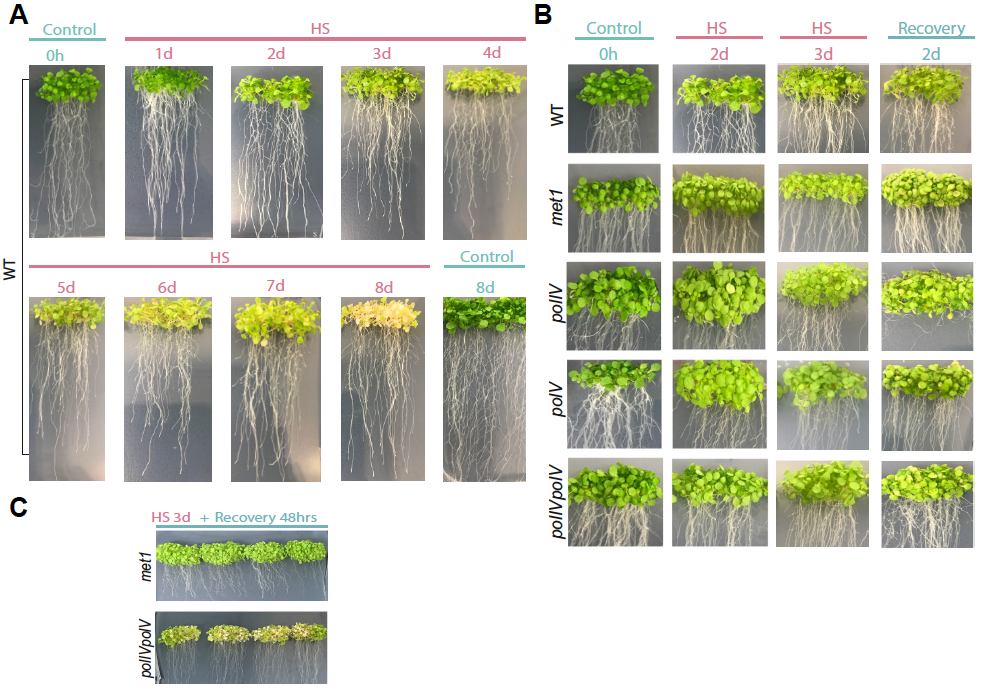

**Fig. S2.** Effect of variable heat stress regimes on health of *Arabidopsis* seedlings. Plant phenotype studied for **A.** Prolonged HS extending up to 8 days using wild type seedlings **B.** Short duration HS applied to both wild and mutant Arabidopsis genotypes**.** Seedlings exposed to heat stress for up to 3d followed by a 2-day recovery phase are shown. **C**.  Phenotype of *met1* and *polIVpolV* mutant seedlings post 3d HS and recovery phase.

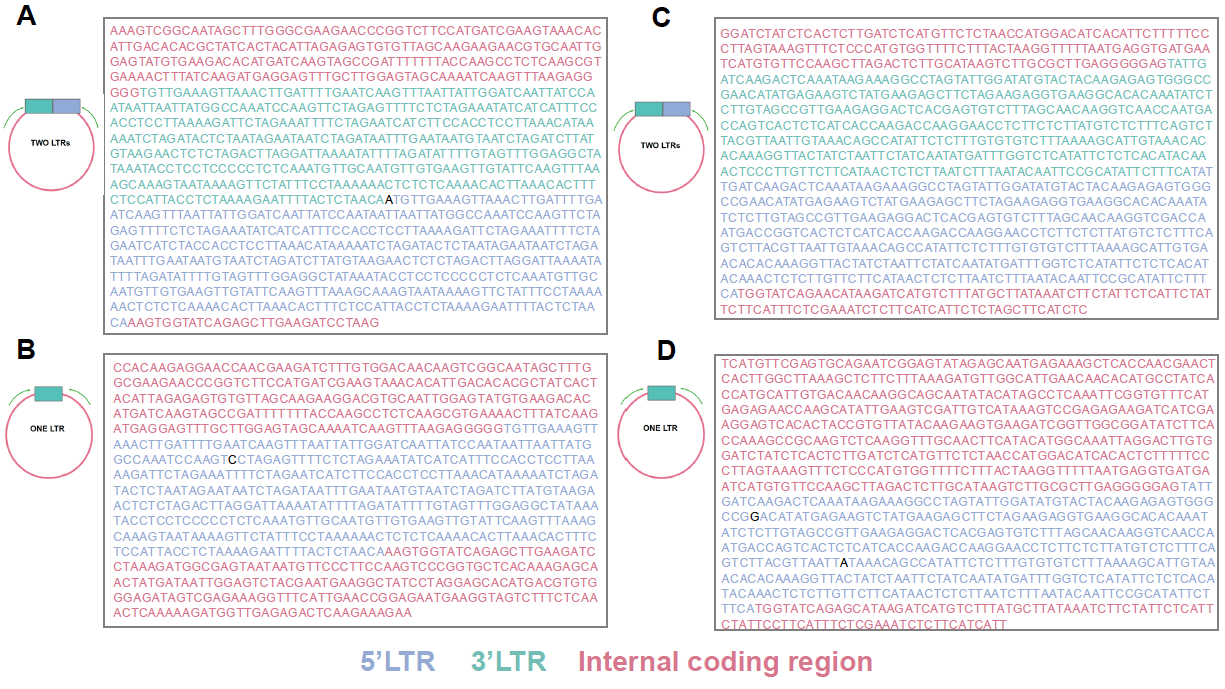

**Fig. S3.** Sequence analysis of two LTR and one-LTR extrachromosomal DNA products of ONSEN (**A, B**) and EVADE (**C, D**). 5’LTR (blue), 3’LTR (green) and Internal coding region (red) are color coded. Variable nucleotides that did not aligned with the reference sequence are shown in black.

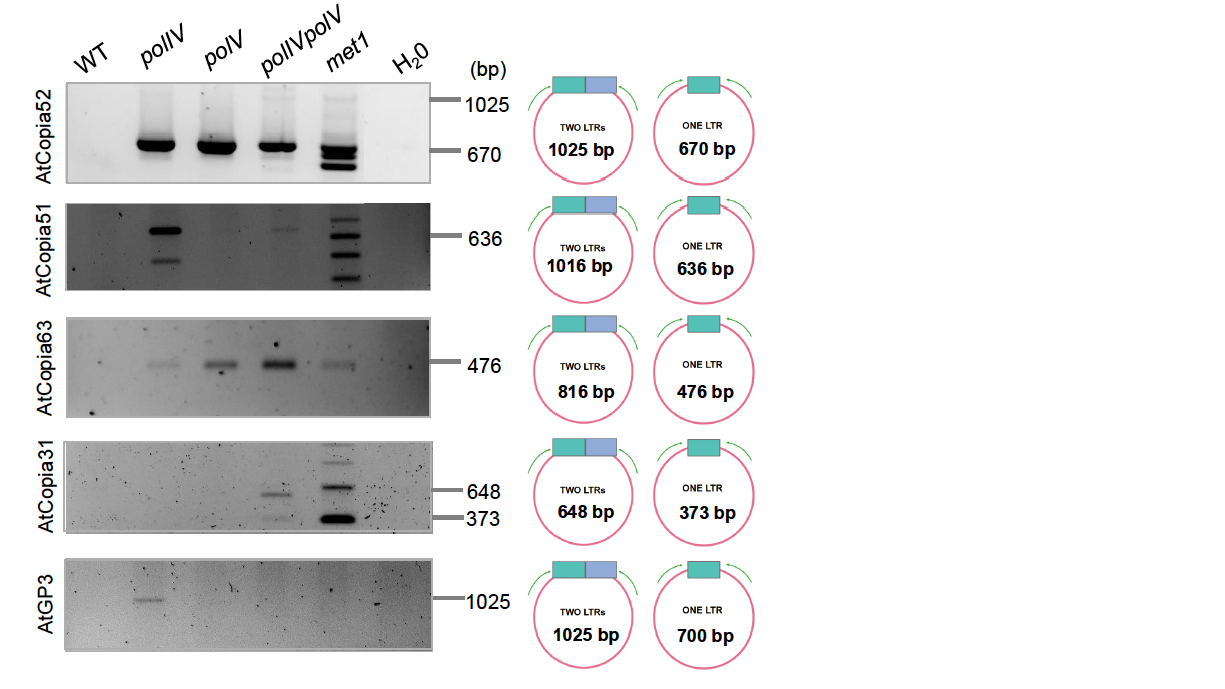

**Fig. S4.** Inverse PCR analysis of eccDNA products of selected epigenetically induced TE families in *A. thaliana* lines. Expected product size for two and one LTR copies have been indicated.

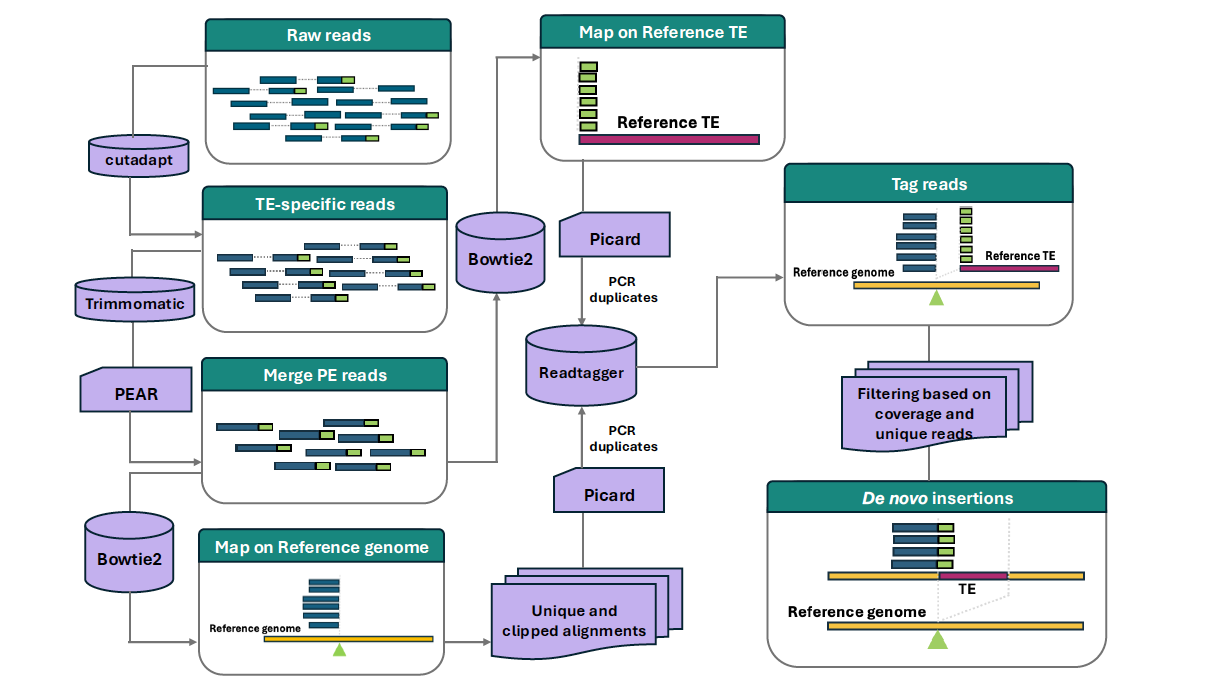

**Fig. S5.** Bioinformatics pipeline “deNOVOEnrich” designed to identify and map *de novo* somatic and non-reference segregating insertions from TEd-seq short-read data. The extremity region of the transposon (from primer site to 5’ end) was used as a bait to harness TE-specific reads from the sequenced library, which were quality filtered. The paired-end reads were merged to create longer contig reads which were mapped separately onto the reference genome and TE. Alignments on the reference genome were filtered to retain uniquely mapped and soft-clipped reads. Soft-clipped reads spanning the insertion site in reference genome wherein their clipped portion showed high homology to the targeted TE were tagged and retained for further analysis. All insertion sites were classified as *de novo* somatic insertions and non-reference segregating insertions based on coverage and PCR duplication information. *De novo* somatic insertions were determined based on their characteristically low coverage, defined as having 5 or fewer supporting reads at the new insertion site with a maximum of four unique reads (defined by distinct stop sites). The non-reference segregating insertions were identified using a coverage threshold of ≥30, with at least five unique reads and a minimum of three of these unique reads supported by at least three PCR duplicates each.

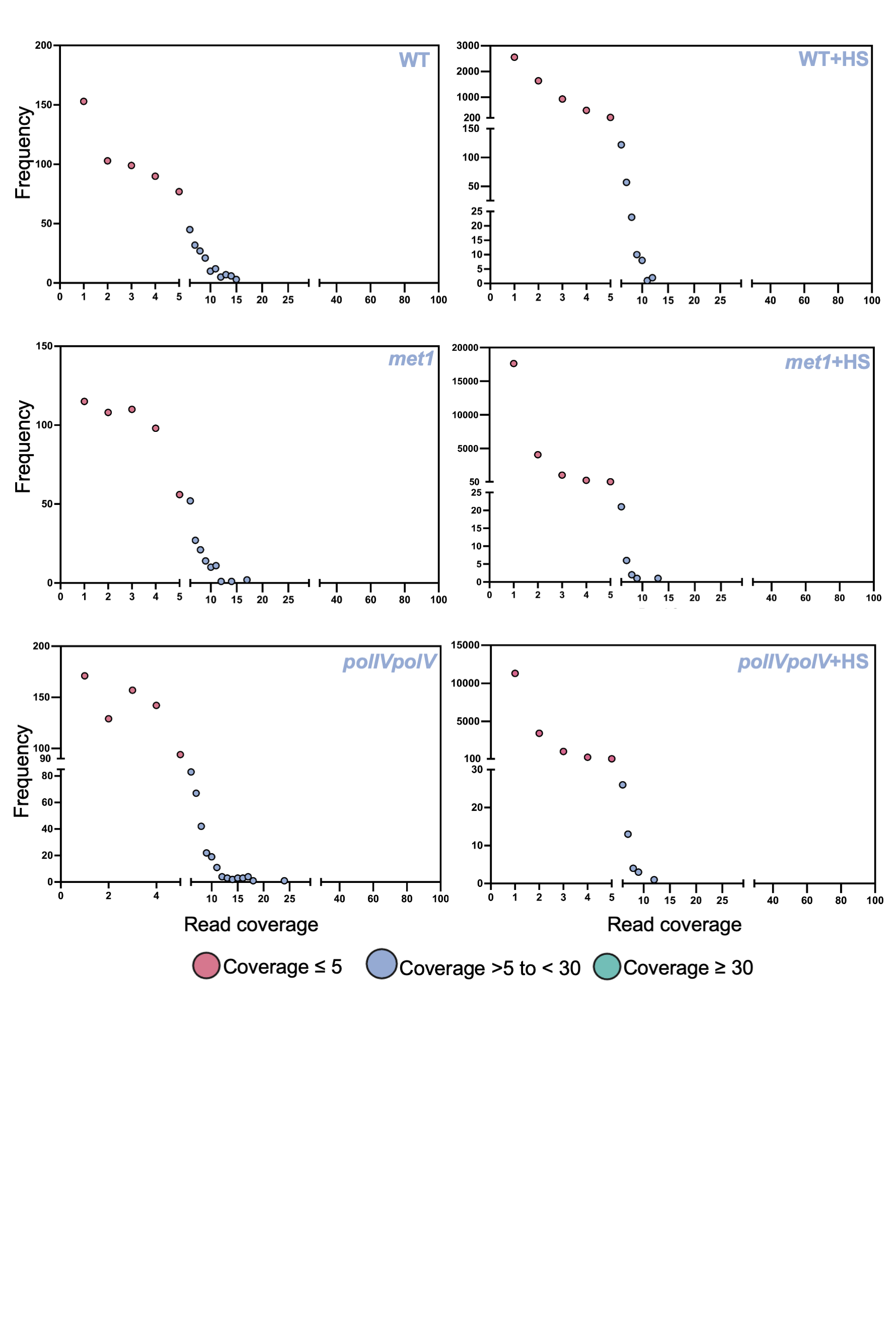

**Fig. S6**. Read coverage support for somatic (<=5 reads) and non-reference fixed (>=30) insertions for ONSEN. Insertions with coverage of >5 and <30 were not further analysed, because of their ambiguous status as somatic or non-reference segregating.

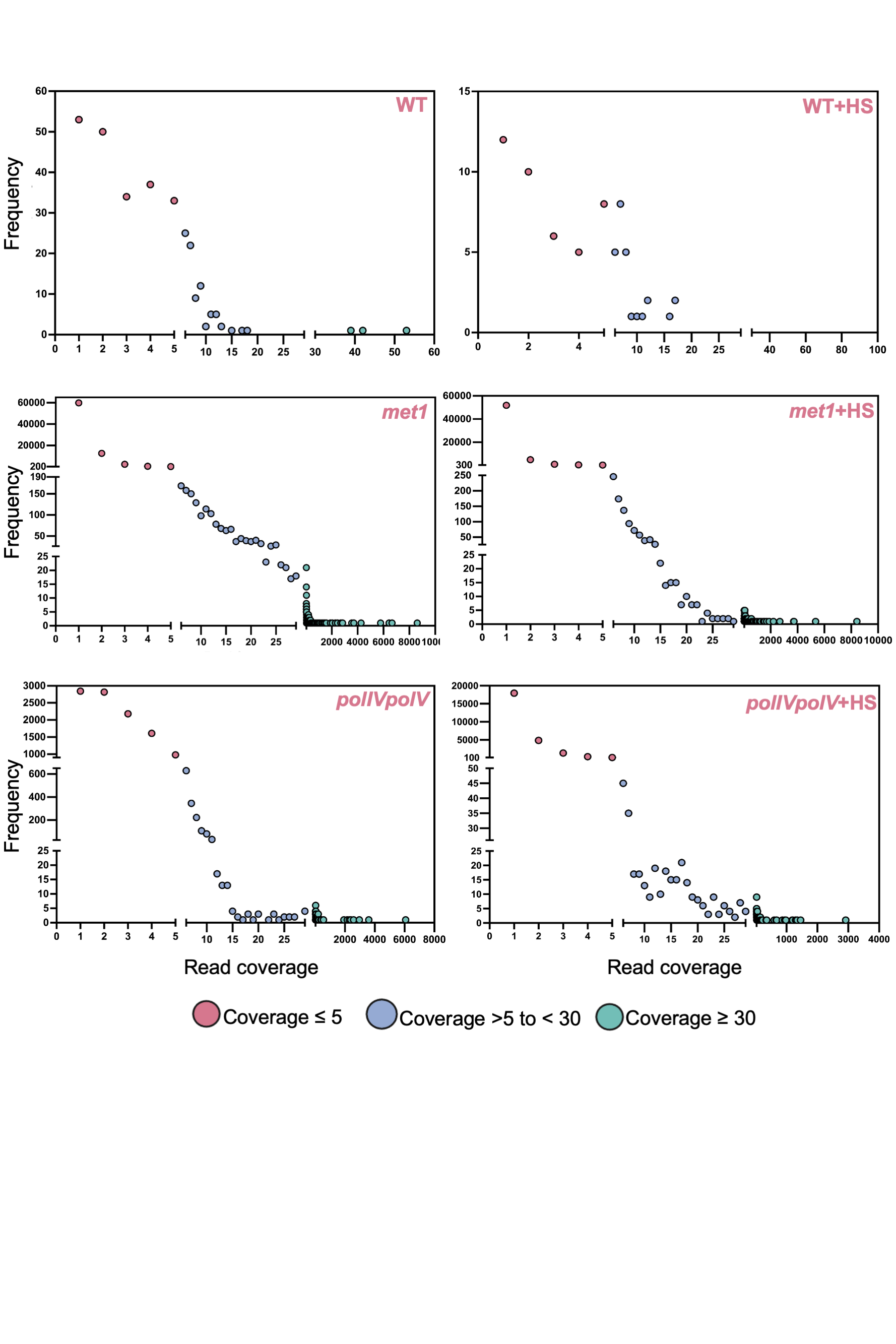

**Fig. S7**. Read coverage support for somatic (<=5 reads) and non-reference fixed (>=30) insertions for EVADE. Insertions with coverage of >5 and <30 were not further analysed, because of their ambiguous status as somatic or non-reference segregating.

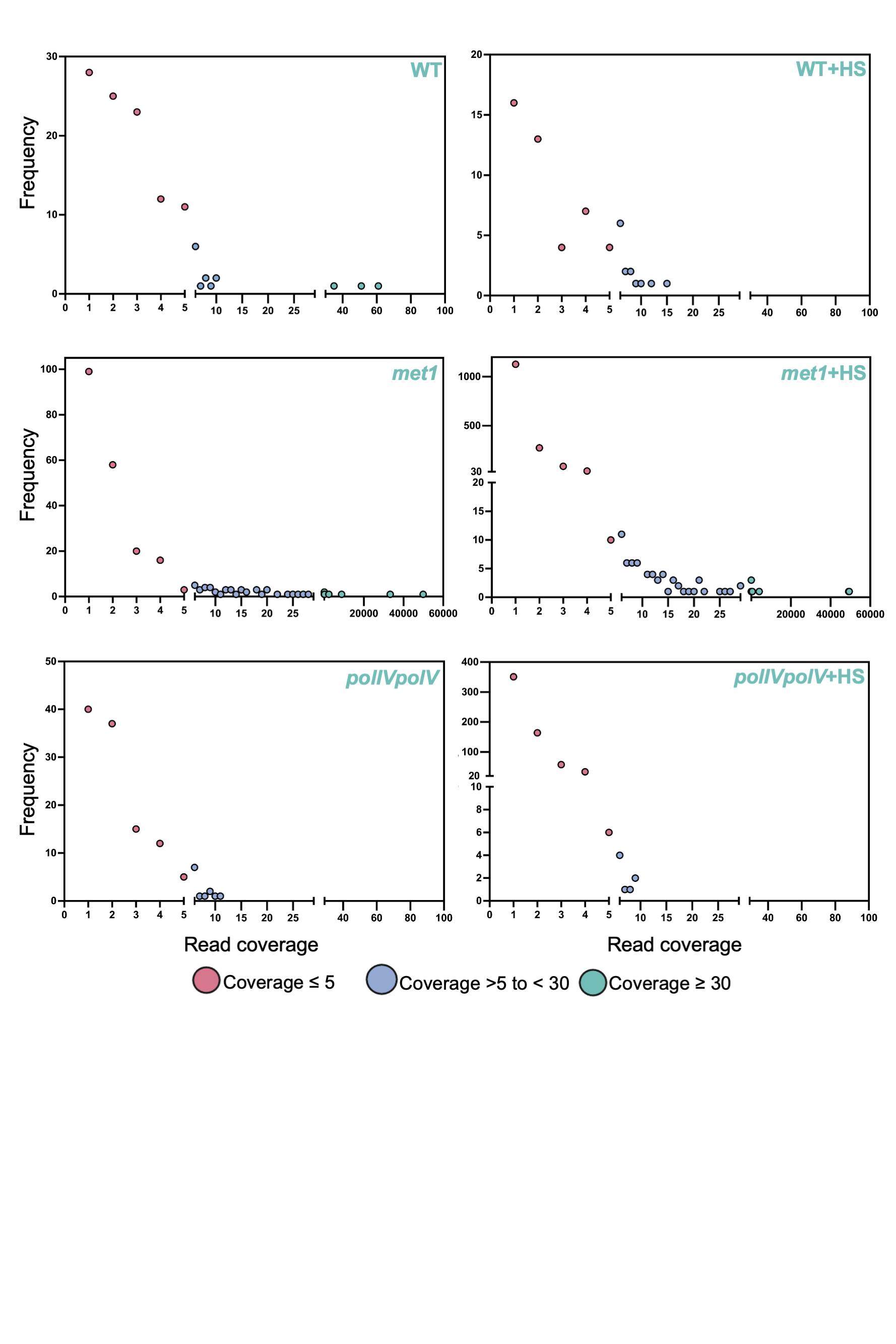

**Fig. S8**. Read coverage support for somatic (<=5 reads) and non-reference fixed (>=30) insertions for AtCOPIA21. Insertions with coverage of >5 and <30 were not further analysed, because of their ambiguous status as somatic or non-reference segregating.

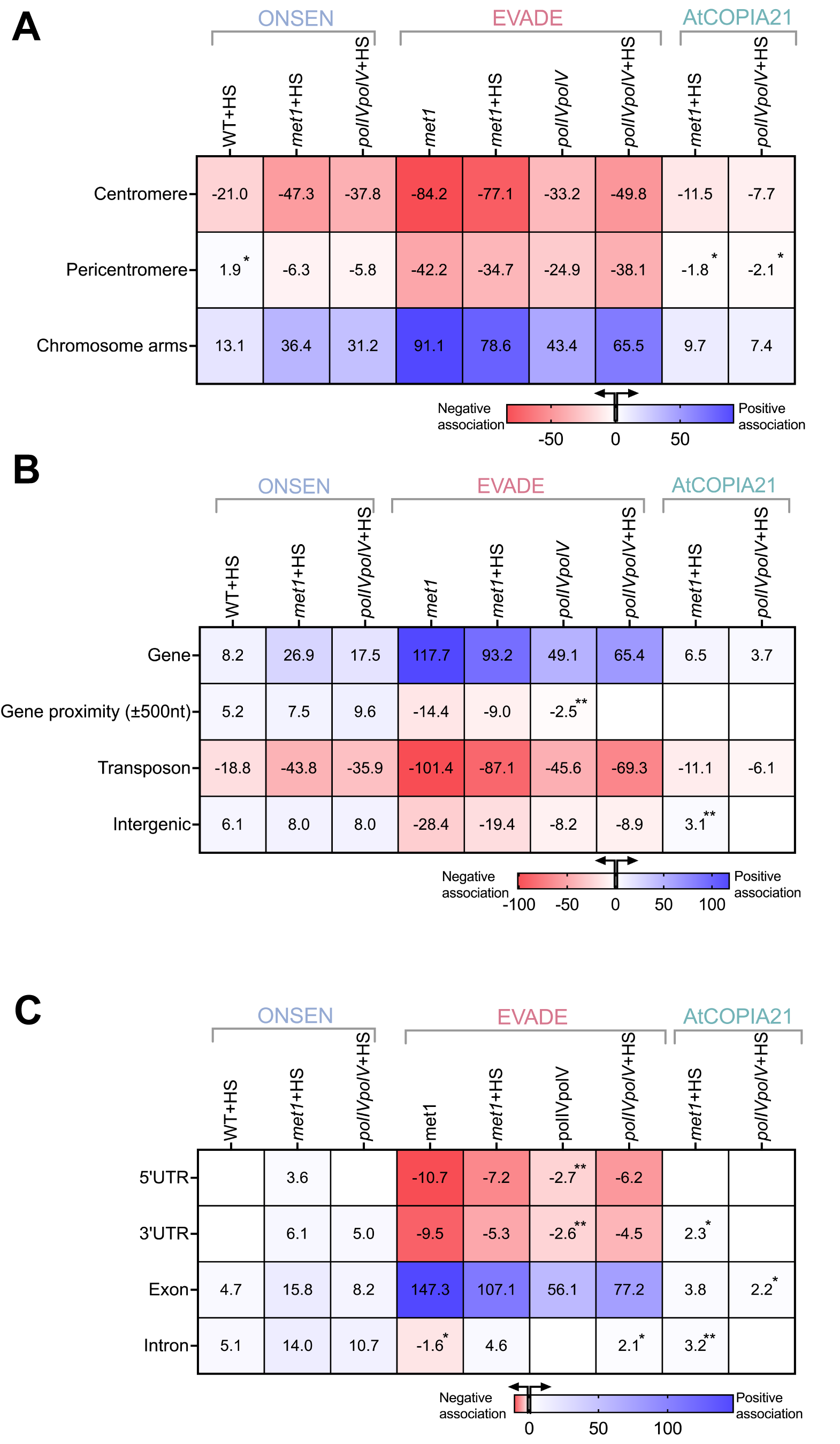

**Fig. S9.** Heatmap depicting permutation Z-scores representing the strength of association between somatic insertions and various genomic features.  **A**. Association with broad chromosomal regions: centromere, pericentromere and chromosomal arms. **B**. Association with genomic contexts: Genes, 500bp windows proximal to genes, reference TEs and intergenic regions **C**. Association with genic components including 5’UTR, 3’UTR, Exons and Introns. Positive associations are shown by blue gradient while negative association represented by red gradient.  Significance is denoted as follows: *p<0.05, **p<0.01 and unless otherwise indicated, all other associations are significant at p<0.001. Empty blocks show samples with non-significant associations.

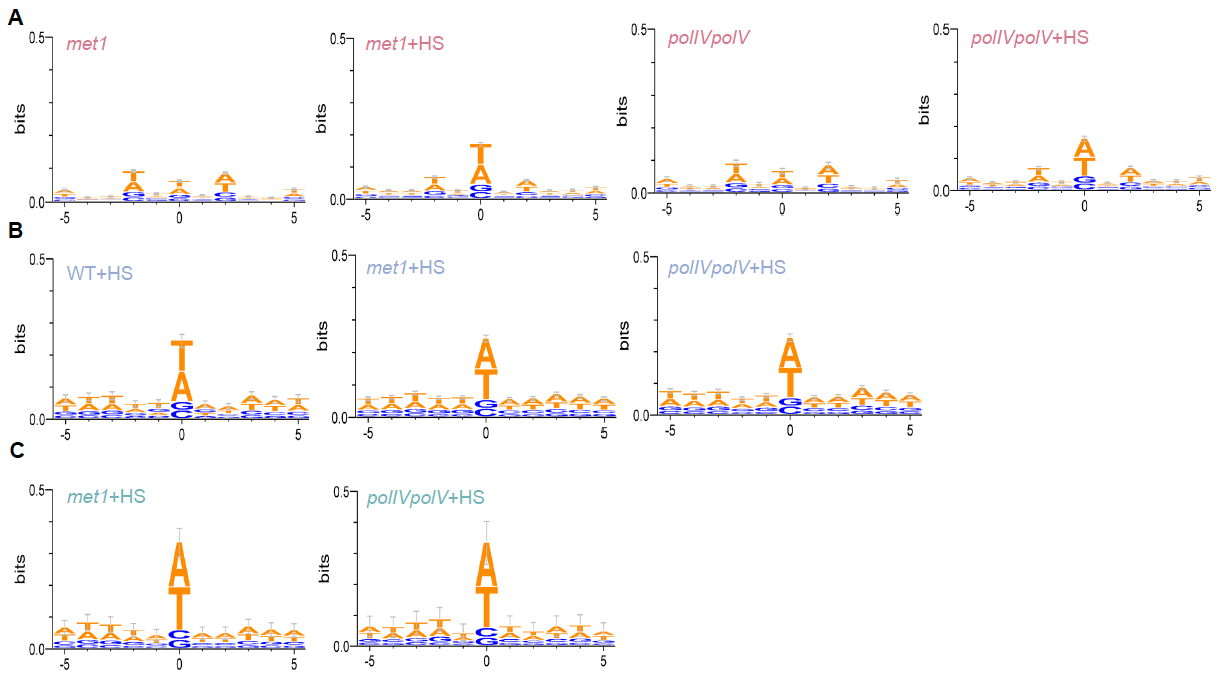

**Fig. S10.** Sequence logo generated using 5bp up- and down-stream of predicted insertion sites of *de novo* somatic transposition events of the three TE families, EVADE (A, red), ONSEN (B, blue) and AtCOPIA21 (C, green).

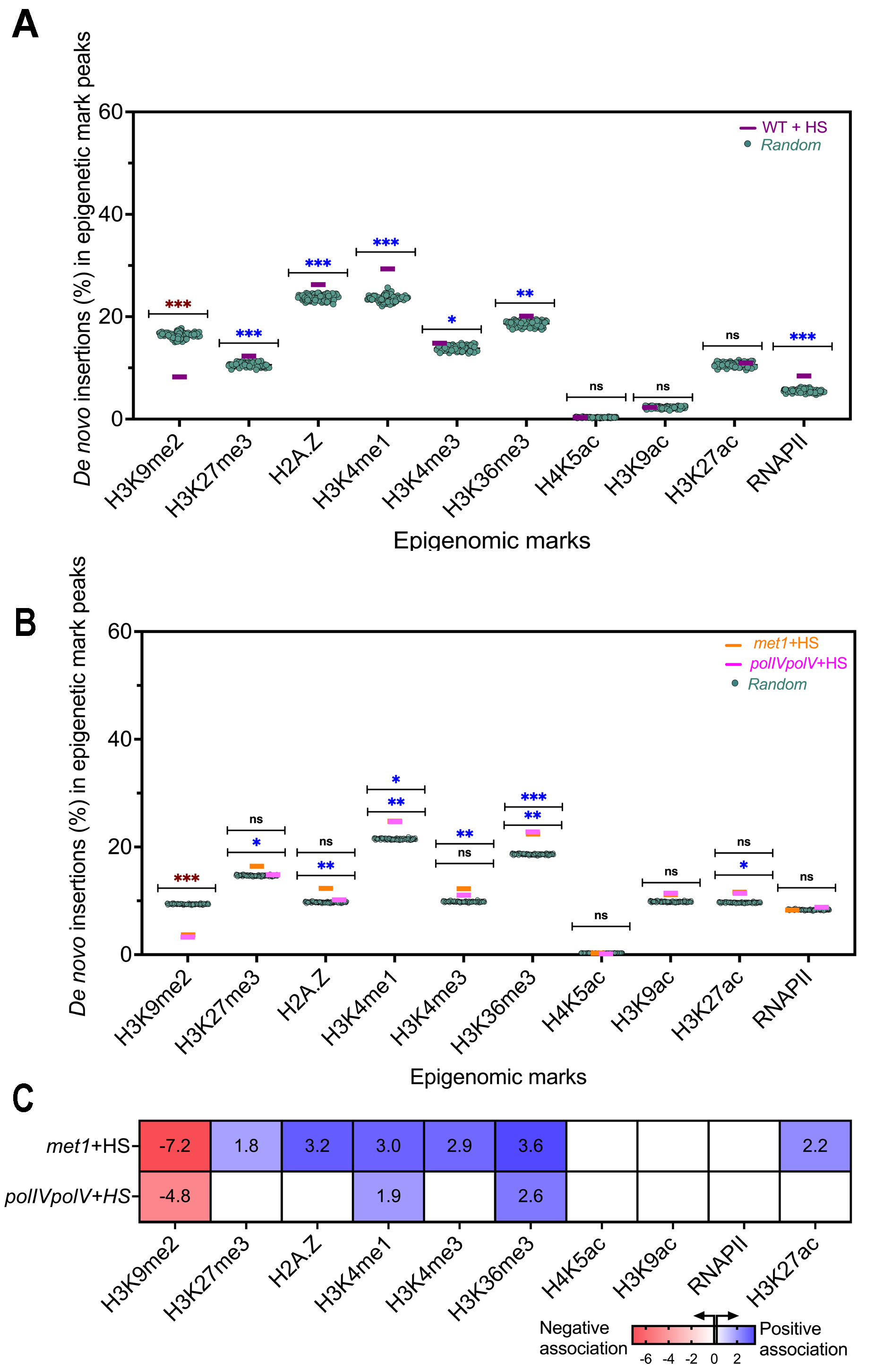

**Fig. S11. A.** Proportion of *de novo* ONSEN insertions in the peak regions of different epigenomic marks in heat-stressed WT tissue. **B.** Proportion of *de novo* insertions of AtCOPIA21 in the peak regions of different epigenomic marks. Colored horizontal lines represent observed values while teal-green dots indicate expected values for the randomized insertions. Permutation test was applied to evaluate level of significance (***p<0.001; **p<0.01; *p<0.05; ns – nonsignificant). Positive association marked by blue asterisk while negative association shown by red asterisk. Two significance bars shows that samples had variable significance values. **C**. Heatmap depicting Z-score representing the strength of association between somatic insertions of AtCOPIA21 and epigenomic marks, computed through permutation tests. Positive association shown by blue gradient while negative association are represented by red gradient. Empty blocks show samples with non-significant associations.

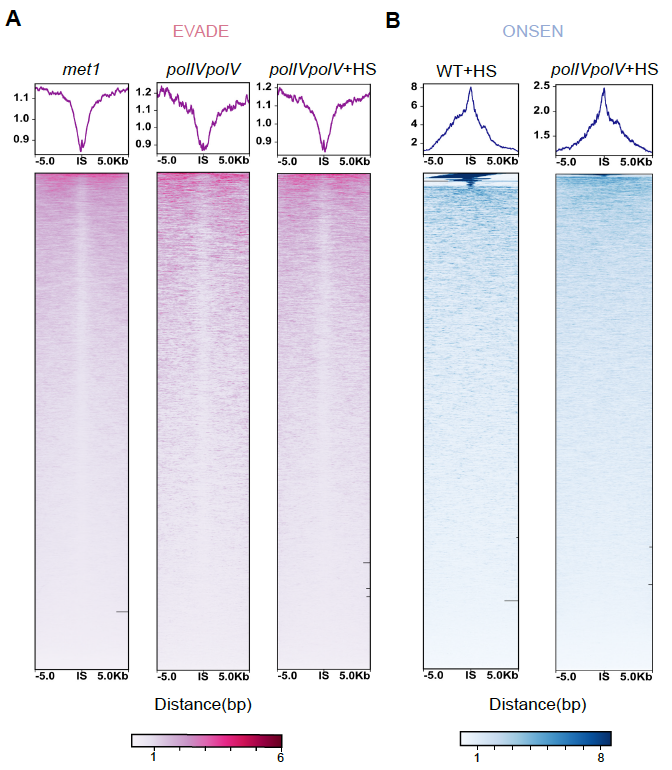

**Fig. S12.** Metagene and heatmap profiles capturing open chromatin at and around (± 5kb flanking region) the insertion sites (IS) of EVADE (A) and ONSEN (B) integration events in different genetic backgrounds and treatments, generated using ATACseq data.

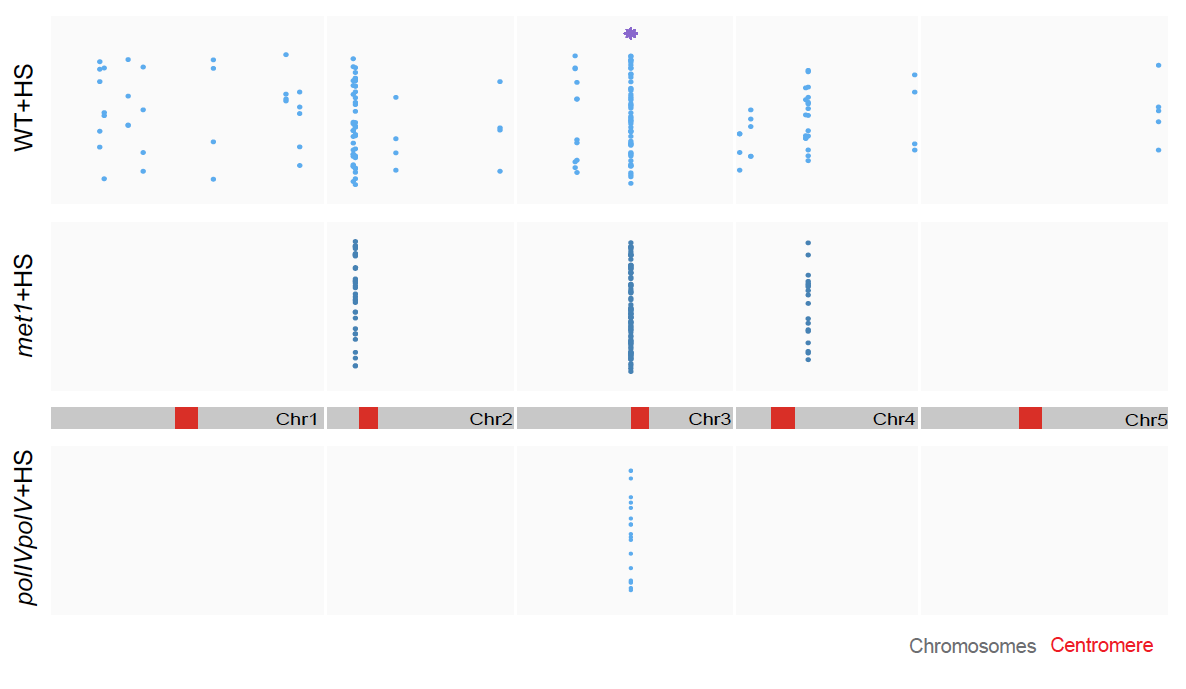

**Fig. S13.** Distribution of insertion hotspots of ONSEN over Arabidopsis genome. Each dot represents an independent insertion event in a 10kb hotspot window. Hotspots shared between the samples have been highlighted by asterisk (purple).

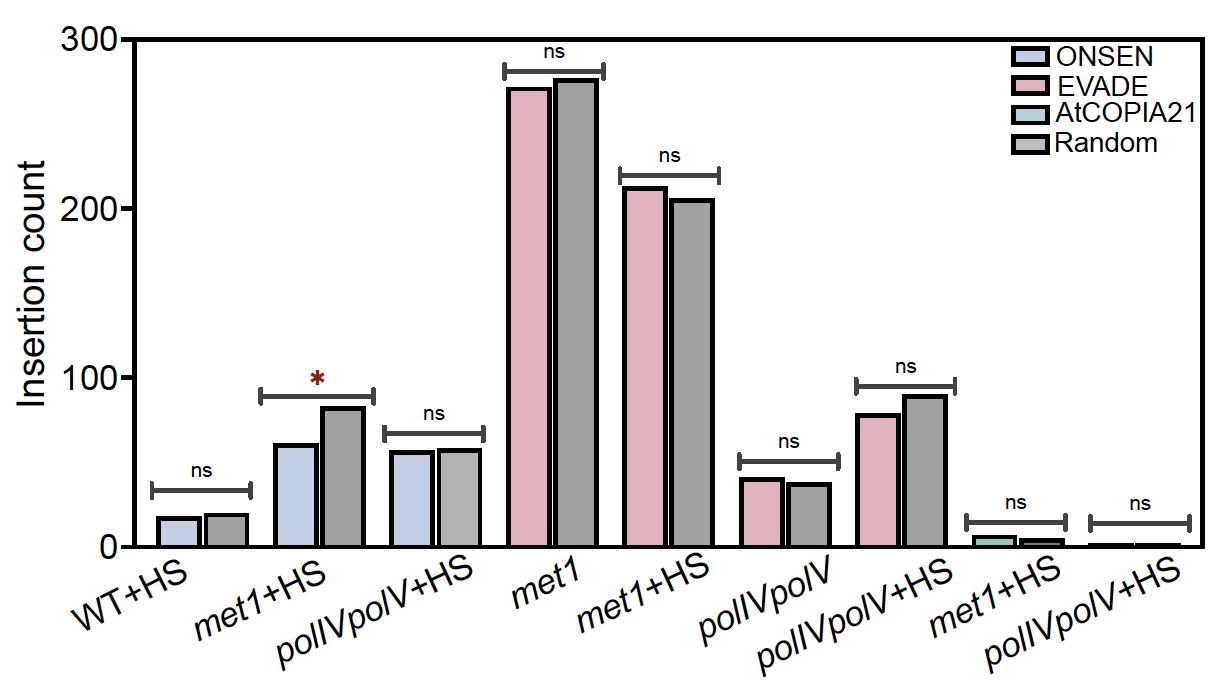

**Fig. S14.** *De novo* somatic insertions of the three TE families overlapping with KNOT regions. Permutation test was applied to evaluate level of significance (*p<0.05; ns - non-significant). Red asterisk indicate negative association signifying depletion of insertions in KNOT regions.

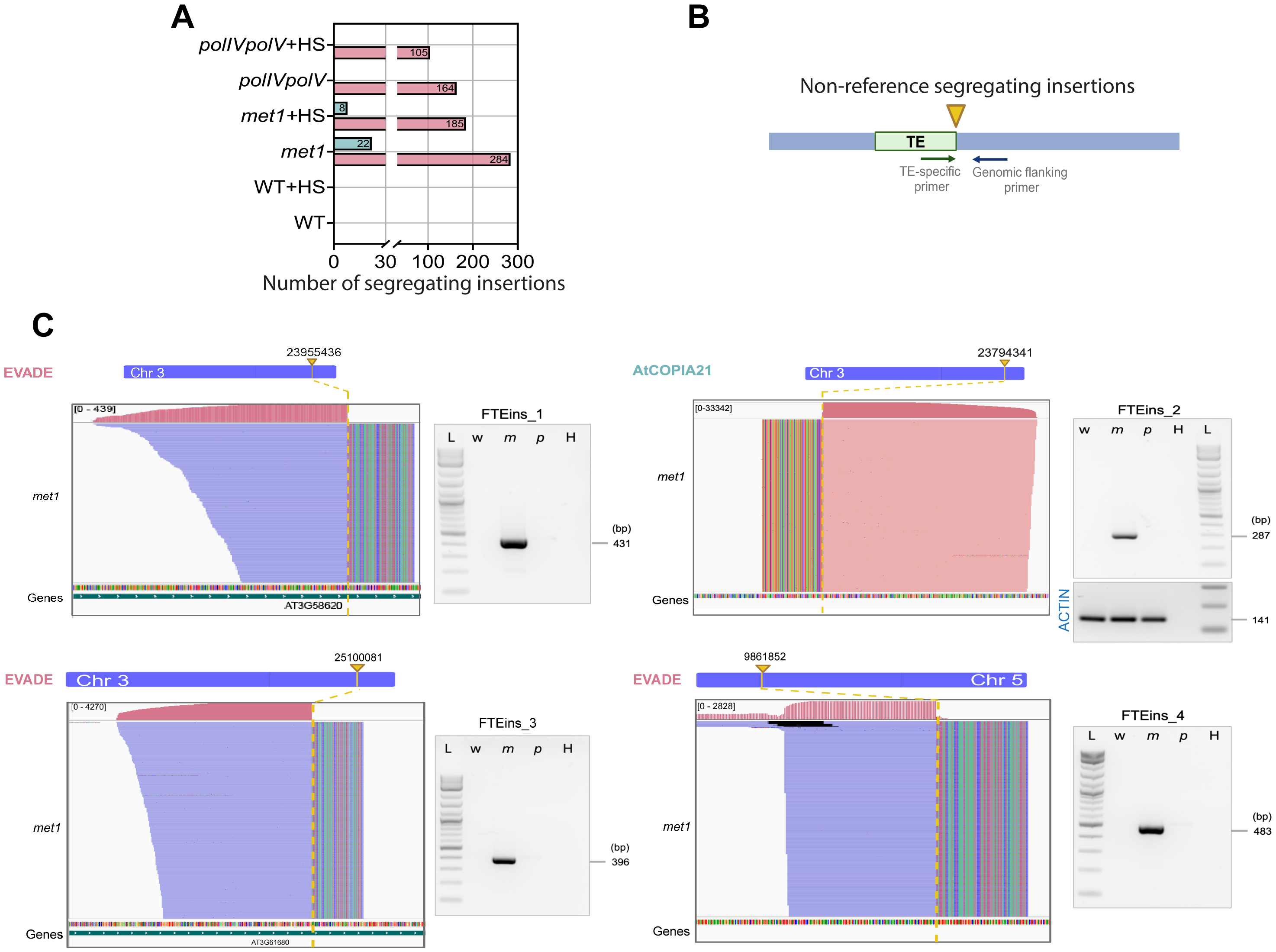

**Fig. S15. A.** Number of segregating TE insertions for EVADE and AtCOPIA21 in wild-type (WT), *met1* and *polIVpolV* with and without heat-stress. **B.** Primer positions (one extending from TE end and other from flanking genomic regions) used for validating segregating transposon insertions. New TE insertion site in the genome has been indicated by inverted yellow arrow. **C.** IGV plots showing position and read coverage at the insertion site of four non-reference segregating TE insertion sites of EVADE and AtCOPIA21 (FTEins_1-4). The rainbow portion of the alignment symbolic for soft-clipped bases corresponds to the TE sequence. Gel profiles for integration site detection by PCR is shown on the right with their expected size indicated. w, wild type; m, *met1*; p, *polIVpolV*; H, water (negative control); L, 1kb plus DNA ladder (NEB).  Amplification from Actin used as control.
