## Additional File 2 for "Somatic mobility of transposons is explosive and shaped by distinct integration biases in *Arabidopsis thaliana*"

**Supplementary Tables**

**Table S1: Plant material used**

| Genotype background | Allele | Functional relevance |
| --- | --- | --- |
| Columbia-0 (Col-0) | Wild type | - |
| *met1* | *ddm2-1* | DNA methyltransferase mediating CpG methylation; loss of function allele |
| *polIV* | *SALK_128428* | Plant-specific RNA polymerase crucial for upstream phase of canonical RdDM pathway |
| *polV* | *SALK_017795* | Plant-specific RNA polymerase critical for downstream phase of both canonical and noncanonical RdDM pathways |
| *polIVpolV* | *nrpd1A-4/nrpe1-11* | Double mutant impairing canonical RdDM pathway and affecting non-canonical pathways |

**Table S2: Genomic features of the analysed TE families**

| TE family | # of intact elements | Total length of intact elements (bp) | # of TE fragments | Total length of fragments (bp) | Total length (bp) | % of genome |
| --- | --- | --- | --- | --- | --- | --- |
| AtCOPIA78 | 11 | 54570 | 14 | 6557 | 61127 | 0.0465 |
| AtCOPIA93 | 4 | 21380 | 7 | 3693 | 25073 | 0.0191 |
| AtCOPIA21 | 1 | 4751 | 5 | 3066 | 7817 | 0.0059 |
| AtCOPIA52 | 1 | 4946 | 4 | 3344 | 8290 | 0.0063 |
| AtCOPIA51 | 2 | 9940 | 5 | 5031 | 14971 | 0.0114 |
| AtCOPIA31 | 3 | 13943 | 4 | 3443 | 17386 | 0.0132 |
| AtCOPIA63 | 4 | 12018 | 12 | 4996 | 17014 | 0.0129 |
| ATGP3 | 3 | 15987 | 39 | 48067 | 64054 | 0.0487 |
| Total | 29 | 137535 | 90 | 78197 | 215732 | 0.1640 |

**Table S3: Details of TEd-seq libraries generated**

| Sample | Condition | Single or Pooled TE library | Raw PE reads |
| --- | --- | --- | --- |
| Col-0 | Control (Unstressed) | Pooled (ONSEN+EVADE+AtCOPIA21) | 37,911,113 |
| *met1* | Control (Unstressed) | Pooled (ONSEN+EVADE+AtCOPIA21) | 38,348,087 |
| *polIVpolV* | Control (Unstressed) | Pooled (ONSEN+EVADE+AtCOPIA21) | 29,236,709 |
| Col-0 | Heat stress 48 hrs | Pooled (ONSEN+EVADE+AtCOPIA21) | 44,574,331 |
| *met1* | Heat stress 48 hrs | Single_ONSEN | 108,643,724 |
| *met1* | Heat stress 48 hrs | Single_EVADE | 34,156,703 |
| *met1* | Heat stress 48 hrs | Single_AtCOPIA21 | 34,150,669 |
| *polIVpolV* | Heat stress 48 hrs | Single_ONSEN | 42,943,809 |
| *polIVpolV* | Heat stress 48 hrs | Single_EVADE | 35,722,140 |
| *polIVpolV* | Heat stress 48 hrs | Single_AtCOPIA21 | 36,019,649 |

**Table S4. Epigenomic datasets from the public repositories used in the present study**

| Epigenomic mark | Genotype | SRA ID | Reference |
| --- | --- | --- | --- |
| H2A.Z | *met1* | SRR22253156, SRR22253157, SRR22253206, SRR22253208 | Zhou et al., 2023 |
| H3K27ac | *met1* | SRR15853563, SRR15853564 | Zhao et al., 2022 |
| H3K27me3 | *met1* | SRR15853557, SRR15853558 | Zhao et al., 2022 |
| H3K4me3 | *met1* | SRR15853555, SRR15853556 | Zhao et al., 2022 |
| H3K4me1 | *met1* | SRR15853559, SRR15853560 | Zhao et al., 2022 |
| H3K9ac | *met1* | SRR15853561, SRR15853562 | Zhao et al., 2022 |
| H3K9me2 | *met1* | SRR15853565, SRR15853566 | Zhao et al., 2022 |
| RNAPII | *met1* | SRR15853567, SRR15853568, SRR15853569, SRR15853570 | Zhao et al., 2022 |
| H2A.Z | Col-0 | SRR11680171, SRR11680170, SRR11680169 | Kim et al., 2023 |
| H3K27ac | Col-0 | SRR15853531, SRR15853532 | Zhao et al., 2022 |
| H3K27me3 | Col-0 | SRR15783051, SRR15783052, SRR15783050 | Kim et al., 2023 |
| H3K4me3 | Col-0 | SRR1635718, SRR1635837, SRR1635390, SRR1635841 | Li et al., 2015 |
| H3K4me1 | Co-l0 | SRR15853528, SRR15853527 | Zhao et al., 2022 |
| H3K9ac | Col-0 | SRR27496336, SRR27496334 | Zhou et al., 2024 |
| H3K9me2 | Col-0 | SRR15853534, SRR15853533, SRR15853537, SRR15853538 | Zhao et al., 2022 |
| RNAPII | Col-0 | SRR5313790, SRR5313791, SRR5313792, SRR5313793 | Liu et al., 2018 |
| H3K36me3* | Col-0 | SRR1635352, SRR1635829 | Li et al., 2015 |
| H4K5ac* | Col-0 | SRR27496335, SRR27496334 | Zhou et al., 2024 |
| ATACseq* | Col-0 | SRR6240778, SRR6240780 | Potter et al., 2018 |

*Col-0 libraries have been used for *met1* and *polIVpolV* integration analysis for these epigenomic marks. ChIP-seq INPUT

**Table S5:** **Primer sequences used in the study for various experimental analysis**

| Experiment | Primer name | Sequence (5' to 3' direction) |
| --- | --- | --- |
| qRT-PCR | 18Sr-F | CGTCCCTGCCCTTTGTACAC |
|  | 18Sr-R | CGAACACTTCACCGGATCATT |
|  | qRT-ONSEN_F | CCACAAGAGGAACCAACGAA |
|  | qRT-ONSEN_R | TTCGATCATGGAAGACCGG |
|  | qRT-EVADE_F | GATAGAGGAGATAGAAGATCTACAACTGG |
|  | qRT-EVADE_R | CTCTATACTCCGATTCTGCACTCGAACA |
|  | qRT-AtCopia21F | CGATTGTTGGAGCGAGACTG |
|  | qRT-AtCopia21R | CAACAACAACAACAACGGCC |
|  | qRT-AtCopia31F | ACAACATCAAGGAGCTTACGG |
|  | qRT-AtCopia31R | GCTTTTAACACTCTCTCTTCCAC |
|  | qRT-AtCopia51F | CGCTTCACCATCATCAGGAC |
|  | qRT-AtCopia51R | TCATCAGAGACACGGAGAGC |
|  | qRT-AtCopia63F | CGTGGTTGGTAGATAGCGGA |
|  | qRT-AtCopia63F | GCCTTCAGACACGACCTTTC |
|  | qRT-SISYPHUS_F | CTGTATAGTGTCGAGAGTGTTCTAGATCC |
|  | qRT-SISYPHUS_R | CCGGAAAAGGGTATGGATCTGCCAT |
|  | qRT-ATGP3_F | CATGGTGACTACAAGGTCGCAGGA |
|  | qRT-ATGP3_R | GTTGCTCAGCGACCGTCGTTTCTAAC |
| eccDNA inverse PCR | eccUniF | TGGCATTTGAGTTGTCTCCC |
|  | eccUniR | CCAATATGGATGGCTTGCCT |
|  | ecONSEN_F | CCACAAGAGGAACCAACGAA |
|  | ecONSEN_R | ATCCTTGATAGATTAGACAGAGAGCT |
|  | ecEVADE_F | TTGAAGTGTGTCGCTCTAATGCTGG |
|  | ecEVADE_R | GCACAAACGGACTGATGAATAAAGC |
|  | ecATCOPIA21_F | GAGGAGGAAGACATAGCCGG |
|  | ecATCOPIA21_R | CACAAACCACAGTCCCGG |
|  | ecATCOPIA31_R | TCACCGACTCCTTTTCACTCA |
|  | ecATCOPIA31_F | AGTGTGTTTTCGGTTTAAGGGA |
|  | ecATCOPIA51_R | CGAGTCTGAGCTGGAGGAGTGAC |
|  | ecATCOPIA51_F | ACTCGGCGTGTCTGCGTCACCGGTCTCA |
|  | ecATCOPIA63_R | TTGCTGCATTGTTGTCTCCA |
|  | ecATCOPIA63_F | TTGCTCAATCAACGACCGAG |
|  | ecSISYPHUS_R | CGACGGAGGTGTCTCCTGTGACGAC |
|  | ecSISYPHUS_F | ACCGCTACCAAGACGAGCCTTCA |
|  | ecATGP3_F | CTGATGACGAGCATCACGATATGGATC |
|  | ecATGP3_R | AGTGCATCCTGTGTTCCATGTTTCGCA |
| Segregating insertion validation | EVD_3LTR_1F | ACAGCCATATTCTCTTTGTGTGT |
|  | FTEins_1R | AGAAGAATCAGCTCCATTTGCAA |
|  | FTEins_2R | AGACATCTGAAGGCTAAGCTGTA |
|  | FTEins_3R | GTATATTCCTTTTGCTGCCTCGT |
|  | AtCOPIA21_3LTR_1F | AGGGTATACACAACTACTAATGACA |
|  | FTEins_4R | TAACACTCTCTCTCTCGTTTCGA |
|  | Actin_F | TGCCAATCTACGAGGGTTTC |
|  | Actin_R | TTACAATTTCCCGCTCTGCT |

**Table S6: Statistics of WGS libraries**

| Libraries | Total raw PE reads | Genome coverage (X) | Total HQ PE reads | Average coverage breadth (%) | Note |
| --- | --- | --- | --- | --- | --- |
| *met1*_HS48hrs | 52103007 | 116 | 49928287 | 99.91 | Leaf tissue harvested from ~100 pooled *met1* seedlings exposed to heat stress (37°C) for 48hrs |
| *polIVpolV*_HS48hrs | 48595300 | 108 | 47201618 | 99.9 | Leaf tissue harvested from ~100 pooled *polIVpolV* seedlings exposed to heat stress (37°C) for 48hrs |

**Table S7: Details of adapter, primer sequences and cycling conditions used for TEd-seq library preparation**

| Library Step | Primer ID | Sequence | Cycling conditions^#^ | Note |
| --- | --- | --- | --- | --- |
| Adapter ligation | P7_adapter_up | GTGACTGGAGTTCAGACGTGTGCTCTTCCGATC*T | NA | *phosphorothioate bond |
|  | P7_adapter_down | pGATCGGAAGAGCATC** | NA | **dideoxy-C ; p: phosphorylation |
| Primary PCR | P7_primer_for | GTGACTGGAGTTCAGACGTG | NA | Adapter-specific primer |
|  | TE78_rev | GGAGGTGGAAATGATGATATTTC | NA | AtCOPIA78 (ONSEN)-specific primer |
|  | TE93_rev | GTGAGTCCTCTTCAACGGCT | NA | AtCOPIA93(EVADE)-specific primer |
|  | TE21_rev | GTGATAAGAGAGTGAATTGTCTTATTATG | NA | AtCOPIA21-specific primer |
| Nested PCR | P7_primer_index4 | CAAGCAGAAGACGGCATACGAGATGCCAATGTGACTGGAGTTCAGACGTG | NA | Illumina P7 primer with index |
|  | P5_TE78_1 | AATGATACGGCGACCACCGAGATCTACACTCTTTCCCTACACGACGCTCTTCCGATCTACTCTAGAACTTGGATTTGGCC | *n*=4; *m*= 8 | Illumina P5 primer with AtCopia78-specific inner primer |
|  | P5_TE78_2 | AATGATACGGCGACCACCGAGATCTACACTCTTTCCCTACACGACGCTCTTCCGATCTTACTCTAGAACTTGGATTTGGCC |  |  |
|  | P5_TE78_3 | AATGATACGGCGACCACCGAGATCTACACTCTTTCCCTACACGACGCTCTTCCGATCTATACTCTAGAACTTGGATTTGGCC |  |  |
|  | P5_TE78_4 | AATGATACGGCGACCACCGAGATCTACACTCTTTCCCTACACGACGCTCTTCCGATCTCGGAACTCTAGAACTTGGATTTGGCC |  |  |
|  | P5_TE78_5 | AATGATACGGCGACCACCGAGATCTACACTCTTTCCCTACACGACGCTCTTCCGATCTGCAGGACTCTAGAACTTGGATTTGGCC |  |  |
|  | P5_TE93_1 | AATGATACGGCGACCACCGAGATCTACACTCTTTCCCTACACGACGCTCTTCCGATCTGCCCACTCTCTTGTAGTACATATC | *n*=4; *m*= 8 | Illumina P5 primer with AtCopia93-specific inner primer |
|  | P5_TE93_2 | AATGATACGGCGACCACCGAGATCTACACTCTTTCCCTACACGACGCTCTTCCGATCTTGCCCACTCTCTTGTAGTACATATC |  |  |
|  | P5_TE93_3 | AATGATACGGCGACCACCGAGATCTACACTCTTTCCCTACACGACGCTCTTCCGATCTATGCCCACTCTCTTGTAGTACATATC |  |  |
|  | P5_TE93_4 | AATGATACGGCGACCACCGAGATCTACACTCTTTCCCTACACGACGCTCTTCCGATCTCATAGCCCACTCTCTTGTAGTACATATC |  |  |
|  | P5_TE93_5 | AATGATACGGCGACCACCGAGATCTACACTCTTTCCCTACACGACGCTCTTCCGATCTGAATTGCCCACTCTCTTGTAGTACATATC |  |  |
|  | P5_TE21_1 | AATGATACGGCGACCACCGAGATCTACACTCTTTCCCTACACGACGCTCTTCCGATCTGTACTTAGTCTAGGGTATACACAACTAC | *n*=6; *m*= 10 | Illumina P5 primer with AtCopia21-specific inner primer |
|  | P5_TE21_2 | AATGATACGGCGACCACCGAGATCTACACTCTTTCCCTACACGACGCTCTTCCGATCTTGTACTTAGTCTAGGGTATACACAACTAC |  |  |
|  | P5_TE21_3 | AATGATACGGCGACCACCGAGATCTACACTCTTTCCCTACACGACGCTCTTCCGATCTATGTACTTAGTCTAGGGTATACACAACTAC |  |  |
|  | P5_TE21_4 | AATGATACGGCGACCACCGAGATCTACACTCTTTCCCTACACGACGCTCTTCCGATCTCACGGTACTTAGTCTAGGGTATACACAACTAC |  |  |
|  | P5_TE21_5 | AATGATACGGCGACCACCGAGATCTACACTCTTTCCCTACACGACGCTCTTCCGATCTGCACTGTACTTAGTCTAGGGTATACACAACTAC |  |  |

\#*n* and *m* represent specific PCR cycle number optimized for each TE family
